# MINGL Quantifies Borders, Gradients, and Heterogeneity in Multicellular Tissue Organization

**DOI:** 10.64898/2026.03.24.713296

**Authors:** Kyra Van Batavia, James Wright, Annette Chen, Yuexi Li, John W. Hickey

**Affiliations:** Department of Biomedical Engineering, Duke University, Durham, NC, 27708, USA; Department of Computer Science, Duke University, Durham, NC, 27708, USA; Department of Mathematics, Duke University, Durham, NC, 27708, USA

**Author notes:** Contributing authors.

## Abstract

Tissues are organized with interacting multicellular organizational units whose interfaces and transitions shape function in health and disease. Current spatial-omics analyses typically assign cells to a single cellular neighborhood—ignoring natural gradients, heterogeneity, and borders. Here we present MINGL (Mixture-based Identification of Neighborhood Gradients with Likelihood estimates), a probabilistic framework that converts existing neighborhood annotations into continuous measures of tissue architecture. MINGL models each cell by multi-membership probabilities across hierarchical organizational units and uses these probabilities to identify enriched cells at interfaces between units, constructs interaction networks across hierarchical scales, quantifies compositional gradient transitions, measures context-specific composition heterogeneity, and provides a starting point for neighborhood resolution. Across multiple spatial-omic datasets spanning melanoma, healthy intestine, and Barrett’s Esophagus progression, MINGL detected innate immune-enriched interfaces at tumor and anatomical interfaces, plasma cell niches linking cellular neighborhoods, distinct regimes of sharp and gradual transitions between organizational states, and disease-associated neighborhood remodeling. By treating neighborhood assignment uncertainty as a biological signal rather than noise, MINGL unifies discrete and continuous representations of tissue organization and makes tissue architecture measurable, comparable, and scalable across biological scales and spatial-omics platforms.

## 1 Introduction

Our bodies are made up of millions of cells that are organized into multicellular organizational units of specific composition and location. These organizational units span across biological scales, coordinate tissue and organ functions, and represent multiple intercellular interactions. When these organizational structures are compromised, destroyed, or altered, disease and organ dysfunction occur—e.g., digestion^1,2^, cancer^3,4^. Recent advances in single-cell, spatial-omics technologies have transformed our ability to characterize and understand these organizational structures in health and disease^5–9^. In tandem, we and others have developed algorithms for identifying locally conserved compositions of cells, termed cellular neighborhoods^10–16^. These powerful approaches have led to the discovery of new immune interactions, disease-associated spatial architecture, and correlations of spatial organization prevalence and interactions with clinical outcomes^17–21^.

However, despite these discoveries and advances in identifying organizational units such as cellular neighborhoods, current analyses remain limited. First, neighborhood algorithms either cluster vectors of cells^14,22–24^ or molecules^25,26^ within a certain radius to group cells in similar niches. This forces cells to a single neighborhood assignment. While useful for comparisons, it does not fully align with our understanding of the biology of cellular neighborhoods, where these can exist either as spatial^27,28^ or developmental^29–31^ gradients of slowly changing organizational structures. Recently, researchers developed GASTON to model continuous transcriptomic contours, revealing molecular gradients across tissue regions^24^. Similarly, MONTAGE infers spatial distributions of functional programs across tissues, identifying regions of low, intermediate, and high functional activity and relating these gradients to underlying cellular communities^32^. However, these approaches primarily characterize gradients at the transcriptomic or functional program level and do not directly identify continuous compositional changes in hierarchical cellular organization. Consequently, identification of cell compositional gradients within or between organizational units remains missing from the spatial analytical toolbox.

Second, while prior algorithms have leveraged inter-neighborhood proximity to identify higher-order associations^11,33^, these algorithms do not characterize the interfaces or borders between them. These biological borders represent critical mixing and regulatory regions that are essential for maintaining higher-order organizational structure. Moreover, these borders are governed by certain cell types usually not found uniform within the organizational units. Yet these cells are currently grouped with broader neighborhood assignments, limiting the ability to identify enrichment or depletion of these important cell types at these borders.

Third, beyond spatial heterogeneity, we know that organizational composition can change according to either disease or inter-personal differences. Yet current algorithms are optimized to separate the most distinct cellular regions, rather than identify changes within organizational units. For example, lymph nodes that are either healthy or metastatic are catalogued as a follicle-based cellular neighborhood^34^. In metastasis, while the composition of the follicle largely remains the same, there are important differences including enrichment of myeloid and fibroblast cell type populations critical to disease progression^34^. However, because of conservation of much of the cell type composition, these changes are missed and lumped within the same neighborhood, limiting our insight into disease- or patient-specific changes that are critical to understanding tissue organization.

Fourth, the selection of the number of neighborhoods to use in these spatial analyses remains largely guess-and-check. This results in many hours of manual curation and user-dependent selection of neighborhoods that can critically impact overall findings. This is especially critical because the depth and complexity of single-cell spatial-omics datasets can vary drastically, so the approach and number used for one study may not apply to the next^9,35^. Although several computational approaches aim to adaptively infer the optimal number of clusters or spatial domains—sometimes accounting for factors such as rare cell populations or cell density—these methods typically produce a single inferred cluster number rather than providing a biologically grounded range of plausible neighborhood numbers^36,37^. While there may not be one “single” correct number, having biologically-grounded recommendations for both a good starting point and range of cellular neighborhoods will help researchers reach conclusions faster and in a more unbiased manner.

Consequently, current spatial analysis methods fail to treat spatial organization simultaneously as discrete and continuous. This limits quantification of organizational features—such as borders and gradients—across hierarchical biological scales. Further, it treats multicellular organization as internally homogeneous units that limit our ability to compare tissue architecture across diverse contexts and conditions. Moreover, the current framework does not provide any explicit representation of uncertainty, number of categories to use, or degree of organizational mixing. Thus, intermediate cellular states and biologically meaningful transitions remain difficult to quantify or compare across tissues, disease states, and biological scales.

To address these computational and conceptual gaps in spatial biology, we developed Mixture-based Identification of Neighborhood Gradients with Likelihood estimates (MINGL). Rather than assigning cells to a single spatial identity (**Fig. 1a**), MINGL treats tissue organization as a probabilistic landscape where borders, gradients, and heterogeneity emerge from uncertainty in organization unit membership (**Fig. 1b**). MINGL leverages uncertainty in organization unit assignments (**Fig. 1c**) to define continuous, interpretable metrics of tissue architecture (**Fig. 1d**). This framework enables unsupervised identification of border cells and organizational unit interaction networks, quantitative measurements of transition gradients and their steepness, direct assessment of architectural heterogeneity across conditions, and a biologically relevant recommendation of cluster number for neighborhood analysis (**Fig. 1e**). By leveraging both discrete and continuous representations of spatial organization, MINGL provides a scalable and generalizable framework for understanding and comparing tissue architecture across diverse biological contexts. MINGL advances spatial analysis from descriptions of multicellular organizational structure, composition, and prevalence, toward quantitative understanding of how multicellular organization is coordinated, conserved, and disrupted at interfaces between hierarchical units.

**Figure 1:**
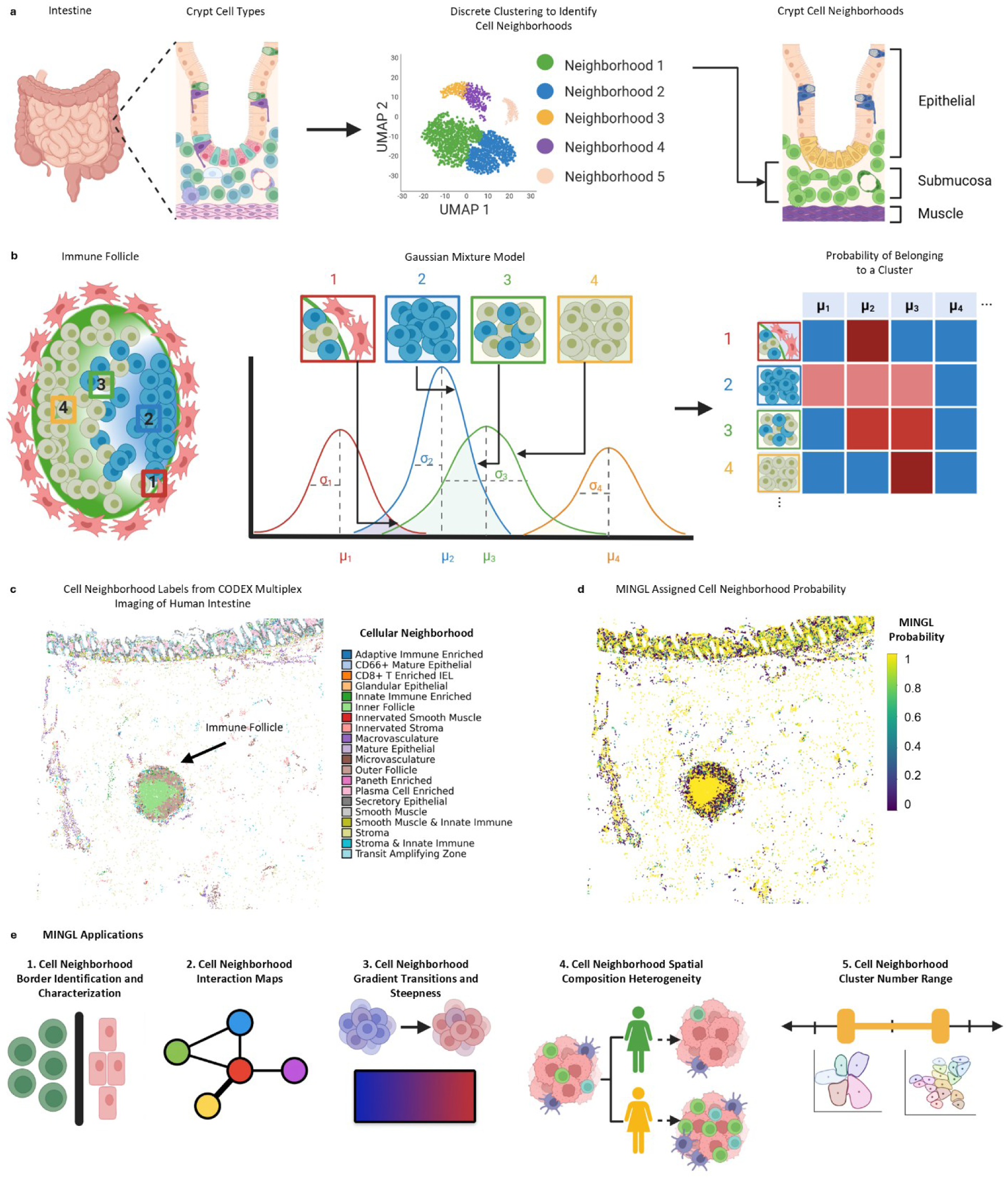
Mixture-based Identification of Neighborhood Gradients with Likelihood estimates (MINGL). **a)** Classical spatial-omics analysis methods rely on discrete clustering of single cells from tissues to define cellular neighborhoods (CNs) of conserved cell composition or coordinated functions. Here, we propose MINGL, a novel probability-based framework for defining continuous spatial organization membership of single cells. **b)** MINGL is based on a Gaussian mixture model where each centroid describes the average features of all possible defined discrete clusters. Each cell is then compared to these distributions, and the probability of cluster membership is calculated. Spatial maps of individual cells’ **c)** assigned CN and **d)** calculated membership probabilities. **e)** These probability maps enable quantitative analysis of tissue architecture, including: **1)** borders between organizational units, **2)** hierarchical interaction networks across units, **3)** spatial gradients and transition dynamics within or between units, **4)** compositional heterogeneity across disease states, individual patients, or tissue regions, and **5)** biologically-grounded selection of an optimal cluster number range as a starting point for spatial-omics analysis.

## 2 Results

### 2.1 MINGL Quantifies Organizational Borders Across Biological Scales

Borders are where organizational units meet, forming regions of biological mixing, cell cooperation and coordination, and transition from one state to another. These borders are not merely physical; they are also hotspots of signaling and interaction that determine tissue behavior. For example, a known biological transition is at the border of solid tumors where immune and stromal cells encounter malignant cells. The composition and spatial hierarchical organization of this tumor boundary is known to be an important factor for effective anti-tumor therapy outcomes^12^. Despite the known biological importance of tissue architecture borders, current single-cell spatial analysis approaches rely on discrete clustering methods that fail to identify border cells.

To identify areas of transition from one organizational state or neighborhood (e.g., tumor) to another (e.g., adjacent normal), we used the multiple predicted probabilities of assignment from the Gaussian mixture model (GMM). We categorize populations of “border cells” that are defined by having two or more CNs above a certain probability threshold (**Fig. 2a**). A CN membership probability above the set threshold is considered a “positive CN membership.” This mathematical definition of a border cell indicates areas of CN blending, which we hypothesize occur at locations where one CN meets another. This definition also enables identification of the presence of borders that are conserved across tissues, tissue types, and individual human donors.

**Figure 2:**
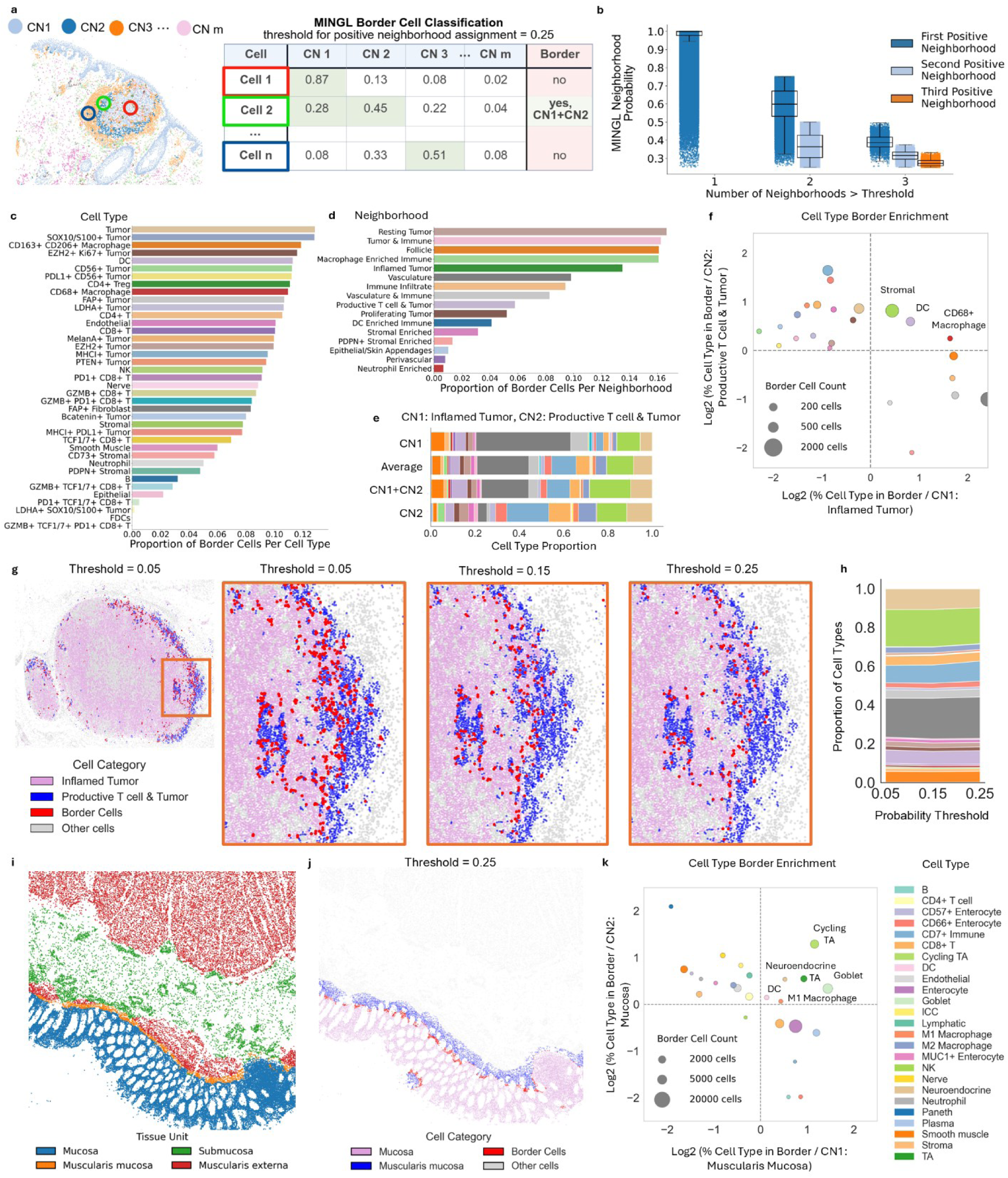
MINGL quantifies organizational borders across biological scales. **a)** Border cells are defined as having positive membership in more than one organizational unit above a certain probability threshold. **b)** Number of positive neighborhoods and probability distributions for cells from the human melanoma spatial dataset. The threshold for positive assignment is 0.25. Proportion of each expert annotated **c)** cell type and **d)** cellular neighborhood (CN) categorized as a border cell between two neighborhoods. **e)** Cell type proportion of cells classified as border cells of *Inflamed Tumor* (CN1) and *Productive T Cell & Tumor* (CN2) CNs compared to non-border cells in each CN and the average of the two populations of non-border cells. Cell type color corresponds to colors in Figure 2c. **f)** Log2-fold enrichment of border cell types in comparison to the *Inflamed Tumor* and *Productive T Cell & Tumor* CNs. Dot color corresponds to cell type (see Figure 2c) and dot size corresponds to the number of cells in the border cell population. **g)** Spatial location of border cells (red) in comparison to the *Inflamed Tumor* (pink) and *Productive T cell & Tumor* (blue) CNs in a selected human melanoma tissue region across different positive neighborhood membership thresholds. Gray cells correspond to cells that are not positive for either selected CN. **h)** Proportion of border cell types across different positive neighborhood membership thresholds. Cell type color corresponds to color in Figure 2c. **i)** Expert annotated tissue unit labels in a representative healthy human intestine tissue region. **j)** Border cell (red) spatial distribution in comparison to the *Mucosa* (pink) and *Muscularis Mucosa* (blue) tissue units. **k)** Log2-fold enrichment of border cell types compared to the *Mucosa* and *Muscularis Mucosa* tissue units. Dot color corresponds to cell type, and dot size corresponds to the number of cells of that cell type in the border cell population.

We applied MINGL at a probability threshold of 0.25 to a human melanoma dataset^12^ to identify border cells in the setting of cancer. At this threshold, the majority of cells (∼91%) were not classified as border cells, ∼9% were classified as border cells, and even fewer (∼0.12%) were above the threshold for three neighborhoods (**Fig. 2b**). This level matches our expectation of compartmentalized cellular neighborhoods where most of the cells will exist within distinct cellular neighborhoods. Moreover, the proportion of border cells did not differ between patient samples, indicating that there are similar levels of border cells across conditions (**Supplemental Fig. 1a**).

To understand whether border cells were located at areas of CN transition, we compared the border cell spatial location to the spatial distribution of CNs. Border cells were most often found in areas of high CN mixing (**Supplemental Fig. 1b, c)**, where high densities of the identified border cells were found spatially enriched within the tumor and tumor-normal border compared to the rest of the tissue (**Supplemental Fig. 1d**). These findings align with our mathematical definition of a border cell as a cell at the intersection of two or more CNs.

Since border cells were restricted to specific tissue locations in our samples, we hypothesized that particular cell types would be enriched at these borders. We found that most tumor cell types had higher proportions of border cells compared to other cell types (**Fig. 2c, Supplemental Fig. 1e**). Of the top 18 cell types with greater than 10% border cell frequency, 11 were tumor cell subtypes—in contrast to healthy epithelial cells where only ∼2% were border cells. These tumor cell subtypes have different functional and phenotypic states (e.g., SOX10+, EZH2+, Ki67+, LDHA+)^38–41^ that are known to have varying outcomes in melanoma patients. Furthermore, observing these tumor cell subtypes intermixing further underscores the spatial tumor cell heterogeneity within tumors^42–44^.

Additionally, immune cell populations were also enriched as border cells, though specific immune cell subtypes were more enriched than others. Both macrophage cell types and CD4+ Tregs and T cells had greater than 10% border cells (**Fig. 2c**). In contrast, B cells and neutrophils were less (3-5%), which suggests that CD4+ T cells are more often found in transition areas of CNs and B cells and neutrophils are more restricted to compartmentalized CNs. Cells that were more frequently found at the intersection of more than one CNs were also enriched within immune and tumor neighborhoods but depleted in normal epithelial neighborhoods (**Supplemental Fig. 1f**). These findings underscore the tumor-normal interface as a unique region of high tumor and immune cell mixing^45–48^, restricted to specific immune and tumor cell types. Specifically, the enrichment of border T cells is consistent with our previous observation of T cell zones at the tumor margin that coordinate anti-tumor immunity^12^.

Because we observed an enrichment of specific CNs within the border cells, we calculated the proportion of cells per CN that were classified as a border cell. Consistent with our cell-type analysis, tumor and immune CNs had the highest proportions of border cells: the *Inflamed Tumor* neighborhood was ∼14% border cells, while the *Tumor & Immune*, *Macrophage Enriched*, and *Follicle* neighborhoods each contained ∼16% border cells (**Fig. 2d, Supplemental Fig. 1g**). In contrast, stromal and epithelial neighborhoods contained a maximum of 4% border cells. These findings indicate that both compartmentalized and consistent neighborhoods such as stromal and epithelial neighborhoods have less frequent border interfaces with other CNs compared to tumor and immune neighborhoods. This indicates that at the cellular neighborhood scale, the tumor and tumor-normal interface is a site of high biological mixing across biological scales contributed to by both diverse tumor phenotypes and diverse immune cell types. This further supports the idea of dedifferentiation or increasing disorganization of cancer in addition to dedifferentiation of the cancer cell itself^49–51^.

We previously found that the neighborhood pair *Inflamed Tumor* and *Productive T cell & Tumor* shared a critical border for anti-tumor responses in melanoma patients^12^. To quantify the biological mixing at this tumor-normal border, the cell type composition of this pair of CNs and their shared border cells were compared. While the composition of border cells more closely mirrored the average cell type composition of the singularly positive CN cells than either individual CN composition (**Fig. 2e**, **Supplemental Fig. 1h**), we also found that stromal cells, dendritic cells, and CD68+ macrophages were more enriched in the border cell population compared to the individual CNs of *Productive T cell & Tumor* and *Inflamed Tumor* (**Fig. 2f**). The enrichment of dendritic cells, CD68+ macrophages, and stromal cells at the interface of *Productive T cell & Tumor* and *Inflamed Tumor* highlight adaptive immune coordination of both antigen presentation or the secretion of inflammatory signals^52,53^, and responding stromal cells that may be involved in inflammation-induced extracellular matrix remodeling or scarring responses^54^.

To determine the dependency of border cell composition and enrichment on the chosen probability threshold, we evaluated a range of threshold probabilities from 0 to 0.25. Border cells were consistently identified across different probability thresholds at the interface where the two CNs spatially meet, with their numbers increasing as the probability threshold decreased (**Fig. 2g**). As expected from the mathematical definition of a border cell, lowering the probability threshold increases the number of cells meeting the criteria for multi-neighborhood membership. Importantly, our analysis showed that border cell type proportions are also consistent across probability thresholds (**Fig. 2h**), suggesting that border cell composition at the tumor-normal border is highly conserved. Our tunable tissue border identification with MINGL suggests that border cells at the tumor-normal interface integrate features of both neighboring tissue architectures, but also certain cell types are enriched at these interfaces and may be important for coordinating separation or transitions between two distinct tissue states.

To further demonstrate MINGL’s identification of borders across diverse tissues and biological scales, we applied MINGL to a human intestine dataset. Unlike the melanoma dataset, this dataset represents healthy tissue rather than disease, involves a different organ system of the intestine instead of skin, and includes annotations spanning larger scale communities and tissue units in addition to cellular neighborhoods (100, 300, 10 nearest neighbors respectively)^11^. Using the neighborhood annotations first at a probability threshold of 0.25, 12% of the cells were classified as border cells similar to the melanoma dataset (**Supplemental Fig. 1i**). Interestingly, again macrophages and CD4+ T were enriched as border cells, whereas specific epithelial subtypes like Paneth cells were less enriched (**Supplemental Fig. 1j**). Paneth cells belong to the *Paneth Enriched* CN that also had a low proportion of border cells, suggesting that this neighborhood is highly compartmentalized (**Supplemental Fig. 1k**).

To understand how tissue borders change at higher biological scales, we applied MINGL using tissue unit level annotations as input to extract tissue unit interactions (**Fig. 2i**). MINGL probabilities of assigned tissue units revealed low probability cells—and thus border cells—at the anatomical borders between the *Mucosa* and *Muscularis Mucosa* and the *Submucosa* and *Muscularis Externa* (**Supplemental Fig. 1l**). Of the border cells identified between the *Mucosa* and *Muscularis Mucosa* (**Fig. 2j**), transit amplifying (TA), cycling TA, goblet, neuroendocrine, DC and M1 macrophage cell types had higher enrichment at the interface compared to the tissue units (**Fig. 2k**). Cycling TA cells were the most strongly enriched at the interface, which is consistent with their spatial localization to the bottom of the intestinal crypt and known to interact with stromal cells and underlying muscularis mucosa in bidirectional feedback that regulates epithelial proliferation, differentiation, and tissue repair^55–57^. The enrichment of DC and M1 macrophages appearing at an interface is interesting as it indicates that these cells may be more involved in borders between organizational structures involving epithelial and stromal cell types.

At tissue unit scale, organizational unit boundaries follow patterns that are well defined in pathology. Here, MINGL defines border cells at the interfaces between the anatomical layers of the intestine. These results emphasize the intestinal mucosa’s high level of cellular interactions^58^ and MINGL’s ability to identify borders at higher biological scales that match anatomical organization patterns. Together, these findings indicate that borders are conserved features of tissue organization across healthy and diseased states and that MINGL provides a robust framework for border cell characterization across hierarchical spatial scales.

MINGL identifies and quantifies border cells at the intersection of more than one organizational unit across biological scales and different probability thresholds. In the melanoma dataset, border cells were consistently enriched at the tumor-normal interface, particularly in tumor and immune cell types and neighborhoods. The identified border cells showed cell type compositions that were intermediate and divergent to the CN pairs they were multiply positive for, revealing potential cell type coordinators of these interfaces. In the healthy intestine dataset, MINGL identified border cells at higher biological scales that matched anatomical organization patterns. By capturing border cells using probability thresholds, MINGL provides a framework for unbiased profiling of tissue boundaries across diverse contexts, providing new insights into regions where cell cooperation and interactions are concentrated.

### 2.2 MINGL Defines Hierarchical Organization Interactions Across Organizational Units

Our work in identifying border cells provided distinct interactions between two or more cellular neighborhoods such as *Inflamed Tumor* and *Productive T cell & Tumor* indicating an ability to detect meaningful interactions between organizational structures. However, how these organizational units interact with each other, the magnitude of these interactions, and the organizational coordination at different biological scales are not well understood. We conjectured that just like you could create a map of a country based on knowing the borders from individual provinces, we could generate an interaction map of a tissue by knowing the borders between organizational “provinces.” We hypothesized that we could quantitatively use the intersection of organizational units from the border cells to identify the interactions of organizational units across biological scales to be able to represent tissue at higher levels of abstraction and extract critical structural relationships in tissues.

To do this, we constructed a network graph using border cell pairs where each node is an organizational unit and the thickness of the edge between two nodes represents the number of cells positive for that pair of organizational units (**Fig. 3a**). We first applied MINGL at a probability threshold of 0.25 in this way to the tissue unit level annotations for the healthy human intestine to understand the prevalence and identities of tissue unit interactions. We observed that the *Muscularis Mucosa* and *Mucosa* have the highest level of interaction (**Fig. 3b**). Almost 100% (0.99) of border cells that were positive for the *Mucosa* were also positive for the *Muscularis Mucosa* (**Supplemental Fig. 2a**). Furthermore, the ordering recapitulated the known spatial layering of the large tissue units in the intestine with the *Mucosa* at the apical surface, followed by the *Muscularis Mucosa*, *Submucosa*, and *Muscularis Externa*. The high levels of *Mucosa* and *Muscularis Mucosa* interaction also reflect the large area present in intestine tissues where these units meet (**Fig. 3c**). Taken together, these findings indicate that MINGL’s mathematical-based network of interactions faithfully reconstruct well understood biological organization at the tissue unit scale and suggest they can be used to construct interaction networks at other biological scales for novel understanding of tissue organization.

**Figure 3:**
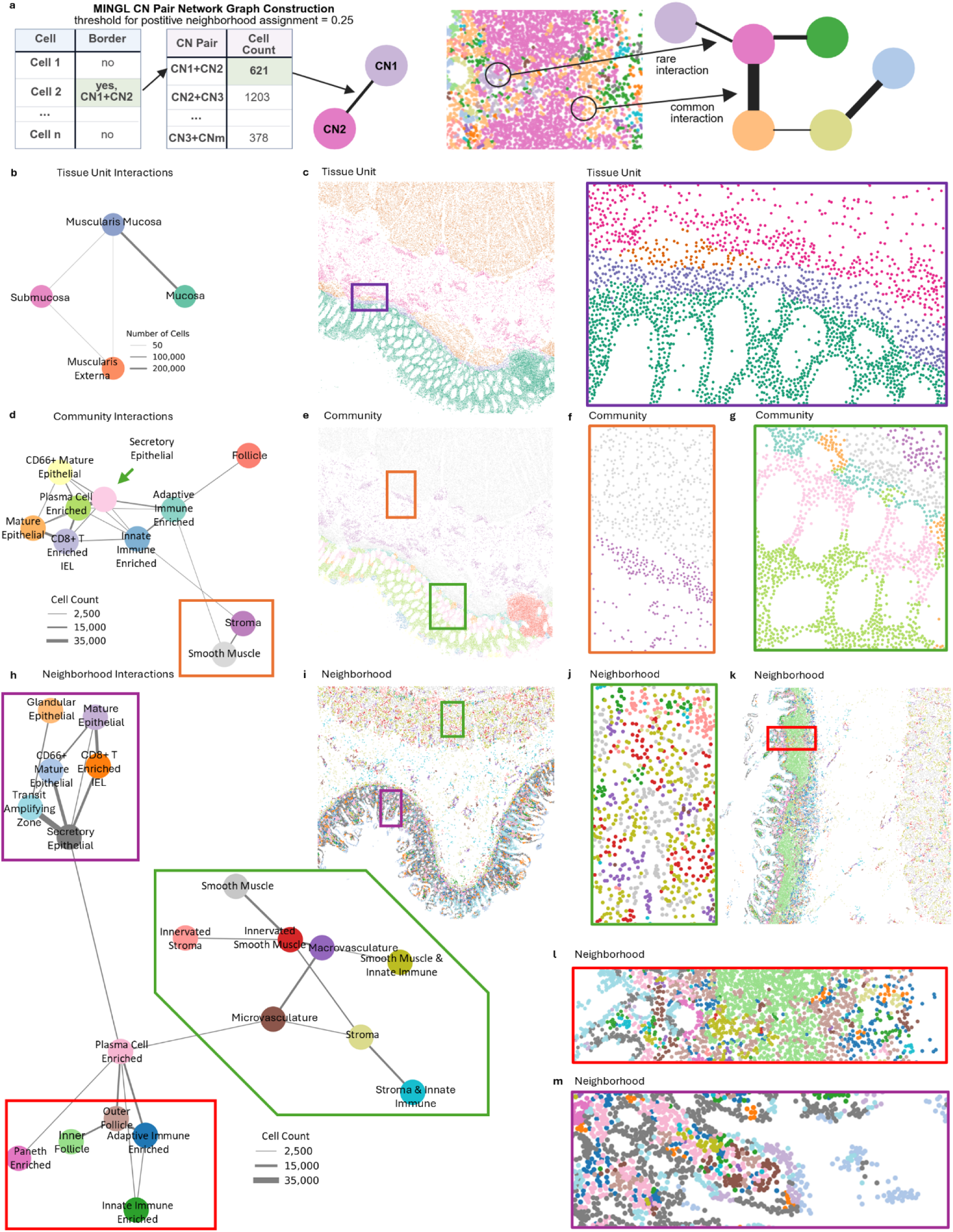
MINGL defines hierarchical organization interactions across organizational units. **a)** Construction of network graphs where each node is an organizational unit and edge thickness corresponds to the number of cells with multi-membership in both nodes. **b)** Tissue unit interactions network graph for a healthy human intestine spatial dataset. **c)** Representative healthy intestine tissue region colored by expert annotated tissue unit labels. Dot color corresponds to annotated nodes in Figure 3b. **d)** Community interactions network graph for a healthy human intestine spatial dataset. **e-g)** Representative healthy intestine tissue region colored by expert annotated community labels. Dot color corresponds to annotated nodes in Figure 3d. **f)** Expanded view of orange box in Figure 3e. *Smooth Muscle* (gray) and *Stroma* (purple) community colocalization. **g)** Expanded view of green box from Figure 3e. *Plasma Cell Enriched* (green)*, Secretory Epithelial* (pink)*, Adaptive Immune Enriched* (teal)*, Mature Epithelial* (orange)*, Smooth Muscle* (gray), and *Stroma* (purple) community colocalization. **h)** Cell neighborhood (CN) interactions network graph for a healthy human intestine spatial dataset. **i)** Representative healthy intestine region colored by expert annotated cellular neighborhood labels. Dot color corresponds to annotated nodes in Figure 3h. **j)** Expanded view of the green box from Figure 3i. Stromal, muscular, and vascular CNs colocalization. **k)** Additional representative healthy intestine region colored by expert annotated cellular neighborhood labels. **l)** Expanded view of the red box from Figure 3k. Immune CNs colocalization. **m)** Expanded view of the purple box from Figure 3i. Epithelial CNs colocalization.

Next, we sought to investigate community-community interactions—a more granular biological scale of the healthy human intestine. The network graph again recapitulated the known organization of the intestine where the *Smooth Muscle* and *Stroma* communities are connected as they correspond largely to the muscularis externa and submucosa of the intestine as we observed with our tissue unit level analysis (**Fig. 3d-f, orange box**). However, the *Stroma* community is connected to a larger network of more defined cell communities that occur within the mucosa of the intestine (**Fig. 3d**). This indicates more diversity of cellular communities within the mucosal area of the intestine.

In particular, the *Stroma* community is directly connected to the *Innate Immune Enriched* community that is also connected to four other cellular communities—with the greatest interaction strength being with the *Adaptive Immune Enriched* community (**Fig. 3d**). If we continue to follow the path of strongest connection in this network graph, *Adaptive Immune Enriched* is connected most strongly to the *Secretory Epithelial* community which is then connected with the *Plasma Cell Enriched* community (**Fig. 3d**). This follows the layering of cellular communities that we observed across the mucosa where we can detect specific borders between the *Secretory Epithelial* and *Plasma Cell Enriched* communities (**Fig. 3e, g, green box**).

Interestingly, we observe the *Plasma Cell Enriched* sharing many edges with diverse communities (**Fig. 3d**). The strongest interaction across the network between the *Secretory Epithelial* and *Plasma Cell Enriched* communities have a co-enrichment score of 0.11 where ∼11% of border cells in the *Plasma Cell Enriched* community are also positive for the *Secretory Epithelial* community (**Supplemental Fig. 2b**). Because the location of this organizational community is conserved to the middle portion of the intestinal crypt (**Fig. 3e, g, green box**), this suggests that the *Plasma Cell Enriched* community is key to the organization of the mucosa. This is somewhat surprising, since the most enriched cell type is plasma cells which are an immune cell and not a structural cell type^59^. Yet there could be a number of reasons that plasma cells are preferentially located in the middle of the crypt. They require protection from epithelial sloughing because they need to be maintained long-term due to their antigen-specificity, yet they also require close access to secrete IgA to the lumen. This is substantiated by the fact that they share the strongest border with *Secretory Epithelial* which have enrichment for cell types such as enterocytes whose main purpose is to mediate transcytosis of IgA into the lumen of the intestine.

We continued to explore this data and these organizational relationships at the most granular level of detail: cellular neighborhoods. The network graph again separated the submucosa and muscularis externa from the mucosa, although there are more subdomains interacting now within these regions such as *Innervated Smooth Muscle* and *Smooth Muscle* (**Fig. 3h, i, j, green box)**.

Again, we observe the more granular *Plasma Cell Enriched* cellular neighborhood as an important crossroads connecting the stromal, epithelial, and largely immune arms of organization (**Fig. 3h**). This can be seen from a section of the healthy intestine where there is a large active immune follicle as represented by the *Outer* and *Inner Follicle* neighborhoods (**Fig. 3h, k, l, red box**). Here the follicular neighborhoods share a border with the *Plasma Cell Enriched* neighborhood—likely a source of plasma cells—that then shares borders with several other epithelial neighborhoods (**Fig. 3l**) and continues as one moves up the intestinal crypt (**Fig. 3h, m, purple box**). This further substantiates the importance of the *Plasma Cell Enriched* neighborhood.

MINGL revealed that neighborhood-neighborhood interactions organize into distinct branches which suggests enriched interactions of neighborhoods within distinct hierarchical organization units. This higher order structure relationship was maintained across both community and tissue unit data. Consequently, by identifying the most enriched border cells, MINGL catalogues the borders of interest between organizational units to focus on. This also provides a useful mathematical formalism in defining higher order multicellular organizational structural interactions across tissues.

### 2.3 MINGL Quantifies Compositional Gradients and Transition Steepness in Organizational Units

Our prior use of MINGL identifies and quantifies borders at defined transitions between two organizational units in tissues. However, the transitions between spatial organizational units are not equivalent; some are stark borders, while others may instead change incrementally. Such spatial gradients represent ordered, continuous transitions within or between organizational units that may be important for organ functions and can be disrupted in disease. Just one example is the immune follicle, where cell types are organized with differential immune cell types that coordinate interactions between cell types—such as between B cells, follicular dendritic cells, and helper T cells at the interface of light and dark zones that enhances the affinity of antibody responses^60–62^. Despite the known biological importance of such spatial gradients, single cell spatial analysis approaches fail to identify transitional zones and quantify how quickly organizational units change and their steepness, limiting our insight into how organizational structures are ordered and maintained in healthy and diseased tissues.

To quantify these transitions, we designed an approach that captures locoregional changes in probabilities of cell neighborhood (CN) assignment at transitions of interest. Based on our prior observations of spatial patterning in assigned cell neighborhood MINGL probabilities (**Fig. 1d**), we hypothesized that these probability gradients could be used to describe cell organization gradients.

First, for a selected CN pair, we first compute a composite probability ratio score for each cell using the log-ratio of the two MINGL probabilities. This transformation sets zero as the mixed-state crossover, weights the ratio by assignment confidence, and enables later quantification of transition steepness. Second, we discretize this continuous score into equal-width bins (e.g., n=5) describing relative similarity (e.g., 80-100 percentile=most like CN1, 0-20 percentile=most like CN2, 40-60 percentile=approximately equal similarity) (**Fig. 4a**) improving interpretability and reducing noise that otherwise produces overly local, non-generalizable patterns. This binning approach preserves the full spread of probability ratio values and maintains the true underlying distribution of cells across bins (e.g., allowing unequal allocation of cells to each bin). Third, we take cells’ nearest N neighbors (e.g., N=20) and count the number of cells in each bin to create a vector of quantities per bin that are clustered and ordered according to probability ratio bins.

**Figure 4:**
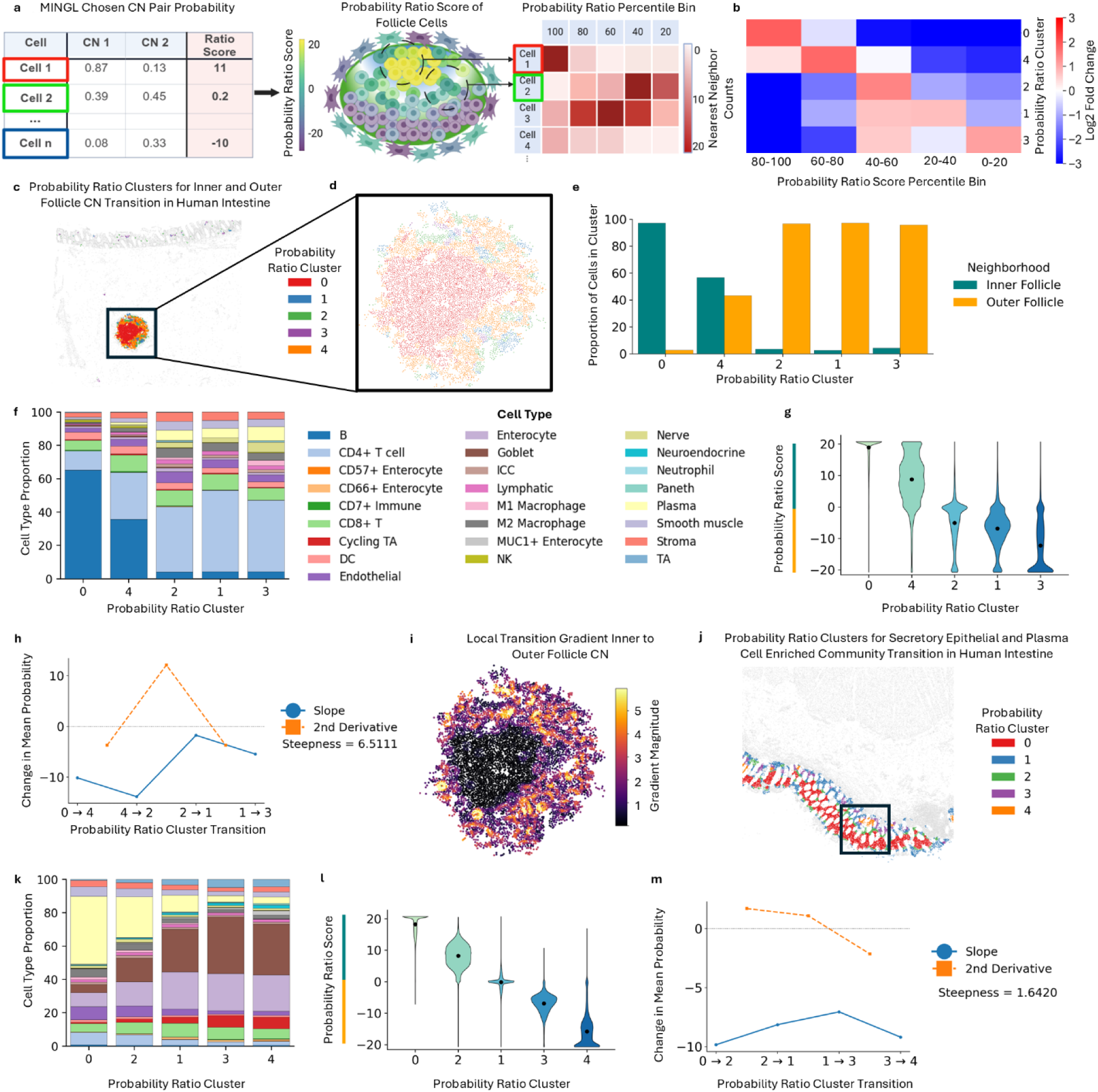
MINGL quantifies compositional gradients and transition steepness in organizational units. **a)** Process for calculating probability ratio scores and obtaining nearest neighbor probability ratio bin vectors. **b)** Log2 fold enrichment of probability ratio percentile across clusters for *Inner Follicle* and *Outer Follicle* cellular neighborhoods (CNs) in healthy human intestine. **c)** Representative region from healthy human intestine dataset. *Inner-* and *Outer Follicle* CN cells colored by their assigned cluster from Figure 4b. Gray cells represent cells not assigned to these CNs and **d)** magnified follicle from region. **e)** Proportion of *Inner-* and *Outer Follicle* CN cells in each cluster. **f)** Proportion of cell types in each cluster. **g)** Violin plot of probability ratio score distributions in each cluster. **h)** Slope and curvature of the change in probability ratio score across ordered clusters with steepness score. **i)** Local gradient magnitudes between the *Inner-* and *Outer Follicle* CNs of a representative healthy intestine immune follicle. **j)** Representative region from healthy human intestine dataset. *Secretory Epithelial* and *Plasma Cell Enriched* communities colored by their assigned clusters. Gray cells represent cells not assigned to these communities. **k)** Proportion of cell types in each cluster from Figure 4j. Color corresponds to cell type from the legend in Figure 4f. **l)** Violin plot of probability ratio score distributions in each cluster from Figure 4j. **m)** Slope and curvature of the change in probability ratio score across ordered clusters with steepness score.

Because MINGL probabilities encode how confidently each cell belongs to its spatial niche, this framework captures uncertainty and spatial mixing during transitions between multicellular organizational units.

We applied this method with MINGL again to the healthy human intestine dataset^11^, and evaluated transitions in cellular neighborhoods in immune follicles. We chose the *Outer Follicle* and *Inner Follicle* CNs because we observed them sharing a border (**Fig. 3h**) and center-to-periphery transitions for follicles have been previously described^62^ yet have not been quantified. Ordering resulting clusters of probability ratios revealed a high-to-low probability ratio enrichment gradient (**Fig. 4b**). Cluster 0 had the largest enrichment for 80-100 percentile probability ratio score cells and was spatially in the center of the follicle (**Fig. 4c**), reflecting the highly conserved and compartmentalized spatial organization of the *Inner Follicle* CN. Cluster 0 was surrounded by concentric bands of clusters with progressively lower enrichment of probability ratio (**Fig. 4d**) and lower proportions of the *Inner Follicle* CN (**Fig. 4e**). Collectively, these results show that unsupervised clustering of MINGL CN probabilities quantitatively defines continuous spatial gradients across immune follicle zones, consistent with known biological transitions.

Next, we examined how individual cell types contribute to these identified spatial organization gradients in immune follicles (**Fig. 4f**). As expected, cluster 0, which is highly enriched for the *Inner Follicle* CN, had the highest proportion of B cells and the proportion of B cells decreased from the center follicle cluster (cluster 0) to the periphery (cluster 3) (**Fig. 4f, Supplemental Fig. 3a**). This result reflects the transition from the B cell enriched dark zone to the surrounding peripheral light zone. More importantly, this indicates that while all of these cell types had previously been identified or binned together as *Inner Follicle* or *Outer Follicle* CN, there exists variance within the neighbor composition of these cells associated with important spatial gradients.

In contrast to B cells, CD4+ T cells increase from the center to the periphery, consistent with their role in B cell support in the follicle light zone. Particularly at cluster 4 we see both intermediate levels of both CNs (**Fig. 4e**) and intermediate levels of B and CD4+ T cells between both extremes pointing to this cluster as a particularly mixed area in the transition (**Fig. 4f**). Additionally, plasma and nerve cells increased from the center to the periphery of the immune follicle (**Supplemental Fig. 3a**). Both changes are aligned with known biology where plasma cells are known to migrate out of the B cell zone after differentiation. Additionally, nerve-associated cells are generally excluded from dense B cell zones, consistent with their enrichment at the boundary^63^. These patterns reveal patterned spatial transitions of cell composition in immune follicle organization that relate to key interactions ongoing in active immune responses.

Our use of clustering probability ratios also enabled us to develop a framework to quantify the rate of transition of organization gradients. Our quantitative metric is derived from the observation of how quickly probability ratio scores change across clusters of probability bins. For example, the maximum (cluster 0) and minimum (cluster 3) represent the range of separation of the two CNs (**Fig. 4g**). This range indicates how distinct the two nodes are in terms of CN composition which would determine the overall slope of change between the two CNs. In between these nodes there could be linear (gradual gradient) or hyperbolic (stark border) trends. Here we observe more of a hyperbolic trend in the change of probability ratio distributions across the clusters indicating a quick gradient from *Inner* to *Outer Follicle* CN (**Fig. 4g**).

To quantify transitions, we calculated the second derivative of the probability ratio score, representing the curvature or rate of change from each cluster to each other (**Fig. 4h**). Two metrics represent how steep the transition is: first, the maximum value of the second derivative specifies how steep one transition is (12 for 4-2 transition); second, the average of the value of all second derivatives represents an average score for how steep transitions are (steepness = 6.51). Because this metric is computed on the same scale, larger scores correspond to faster transitions.

We then also developed a complementary spatial definition of the transition between neighborhoods in space. Our local steepness score estimates how rapidly the probability ratio score changes in the immediate neighborhood surrounding each individual cell. We consider each cell’s closest 20 neighboring cells and use their spatial positions and probability ratio scores to estimate the local rate and direction of change across the tissue. Because this metric is computed at the scale of individual cells, it could be useful for identifying specific portions of the transition area that are more or less steep than another and generates a detailed spatial map of continuous transition intensity and direction across entire tissues. We observe the steepest organization transitions at the spatial interface between the *Inner* and *Outer Follicle* CN (**Fig. 4i**). These spatially steep areas are correlated with cluster 4 which has the highest average local gradient magnitude (**Supplemental Fig. 3b**), and which corresponds well with our derivative analysis that identified the transition from cluster 4 to cluster 2 as the area with the steepest transition (i.e., largest curvature) (**Fig. 4h**).

To compare our gradient analysis to another set of organizational units at another scale, we did the same analysis with the *Plasma Cell Enriched* and *Secretory Epithelial* community MINGL probabilities which was also a highly enriched community-community interaction (**Fig. 3d**). We similarly observed a gradient in probability ratio bin cell enrichment (**Supplemental Fig. 3c**) and a spatial ordering of clusters in the intestine mucosa (**Fig. 4j, Supplemental Fig. 3d**). This ordering reflected change in the abundance of each community, with cluster 0 most enriched with the *Plasma Cell Enriched* community and cluster 4 most enriched for *Secretory Epithelial* (**Supplemental Fig. 3e**).

As expected, the proportion of plasma cells decreased through the transition from *Plasma Cell Enriched* to *Secretory Epithelial* (**Fig. 4k, Supplemental Fig. 3f**), whereas the proportion of secretory cell types of goblet and paneth cells and epithelial cell types of enterocytes and transit amplifying cells increased (**Supplemental Fig. 3g**). Plotting the distributions of cluster associated probability ratio scores indicated a more gradual change than our prior comparison with follicle CNs (**Fig. 4l**). Indeed, calculating the slopes and second derivatives revealed close to 0 values for all transition curvatures, with a maximum value of 2.1 and an average of 1.64 (**Fig. 4m**), aligning with visual interpretation of lower local gradient magnitudes represented in the graph (**Supplemental Fig. 3h**). This indicates that the transition between the *Plasma Cell Enriched* to *Secretory Epithelial* communities is more gradual than that of the immune follicles and demonstrates how this method can be used at other biological scales.

Together, these results underscore how MINGL probabilities encode not only discrete cellular neighborhood identities but also continuous spatial gradients that organize complex tissues. By quantifying gradient steepness, directional transitions, and shifts in cell-type composition, MINGL enables direct measurement of how rapidly organizational states change and where transitions are most spatially concentrated. The ability to distinguish between steep or gradual transitions further enabled quantitative comparison of transitions between biological contexts and biological scales (e.g., *Inner* and *Outer Follicle* CNs versus *Plasma Cell Enriched* and *Secretory Epithelial* communities). By moving beyond boundary detection to measure transition speed, this gradient-based analysis of MINGL provides a framework for systematically interrogating how tissue architectures are organized, maintained, and altered across diverse biological contexts. Importantly, it reframes tissue boundaries from descriptive qualities into quantitatively defined, comparable properties of tissue organization.

### 2.4 MINGL Defines Heterogeneity in Organizational Units Across Conditions and Samples

In our prior use of MINGL, we have demonstrated how we can use variance in the cellular neighbor vectors to define borders and transitions between cellular neighborhoods. Individual multicellular organizations can also vary with differences in development stage, between organs, along a disease trajectory, or between individuals, to name just a few sources of variation.

However, classically the field has not focused on compositional differences and heterogeneity within a distinct organizational structure but instead has focused on differential abundance of organizational structures in comparisons, such as between healthy and diseased tissues. Indeed, there is not a robust quantitative method for measuring heterogeneity in organizational structures across conditions.

To solve this problem and measure heterogeneity in spatial organization of cellular neighborhoods and compare across conditions, we again use MINGL assigned neighborhood probabilities. Here, we compute delta changes in MINGL assigned probabilities between global and group-specific subsets of data (**Fig. 5a**). First, global MINGL probabilities are computed using all data. Next, the dataset is divided into groups for condition-specific (e.g., tissue pathology, developmental stage, disease, region, etc.) MINGL calculations. Finally, the difference between condition-specific probabilities and the global probabilities is calculated to understand how spatial organization composition changes even when proportional differences are not detected at the global level. The direction of the raw delta values describes whether the cellular neighborhood (CN) organization is becoming more separated (positive delta) or more similar (negative delta) to the other CNs in the group compared to global MINGL probability calculations.

**Figure 5:**
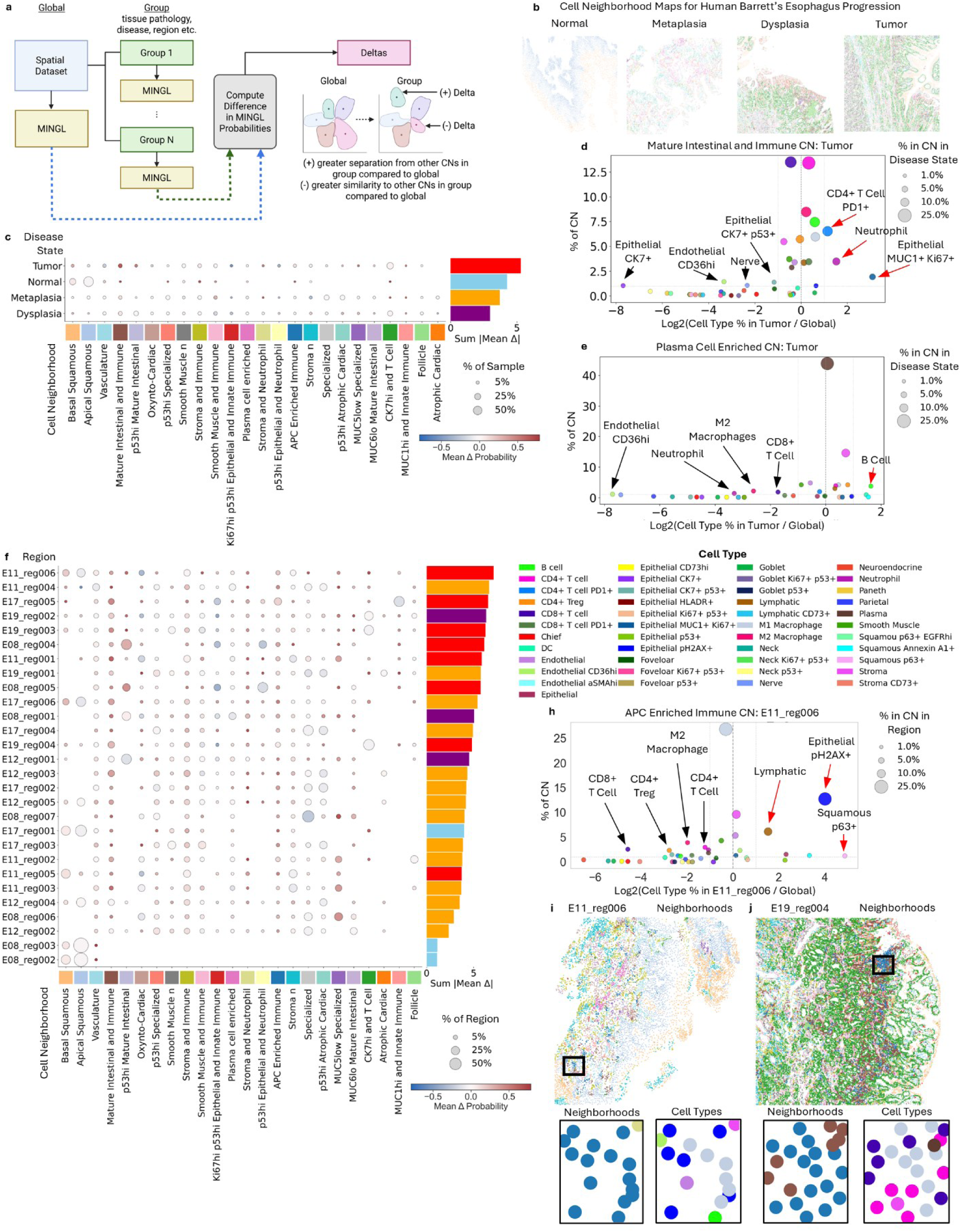
MINGL defines heterogeneity in organizational units across conditions and samples. **a)** Process of calculating delta scores as the difference in MINGL probabilities of cell groupings compared to global. **b)** Representative Barrett’s Esophagus tissue regions of normal, metaplasia, dysplasia, and tumor phenotypes. Dot color corresponds to cell neighborhood (CN) in Figure 5c. **c)** Mean delta scores of each CN across disease state contexts. Dot size corresponds to the proportion of all cells in that context that belong to that CN. Dot color corresponds to the mean delta probability. Sum of the absolute value of mean deltas across all CNs in each context are shown in the bar chart (right). **d-e)** Log2 fold change in **d)** *Mature Intestinal and Immune* CN cell type proportions in tumor disease context compared to global and **e)** *Plasma Cell Enriched* CN cell type proportions in tumor disease context compared to global. For both graphs, dot color corresponds to cell type, and dot size corresponds to the proportion of each cell type in the tumor disease context. Annotated cell types are greater than 1% of the tumor context and greater than 1-fold enriched (red arrow) or depleted (black arrow) compared to global. **f)** Mean delta probability scores of each CN across all Barrett’s Esophagus progression unique regions (n=28). Dot size corresponds to the proportion of CN in each region and dot color corresponds to mean delta score. Bar color (right) corresponds to disease state of unique region (red=tumor, yellow=metaplasia, purple=dysplasia, blue=normal). **g)** Sum of mean delta scores for each patient showing contributions of each unique region. Bar color corresponds to disease state as in Figure 5f. **h)** Log2 fold change in *APC Enriched Immune* CN cell type proportions in specific tissue region E11_reg006 compared to global. Dot color corresponds to cell type in Figure 5e and dot size corresponds to the proportion of each cell type in the specific tissue region. **i)** Barrett’s Esophagus tissue region E11_reg006 colored by expert annotated CN; color legend in colored boxes along x axis of Figure 5f. Extended view of black box region colored by CN (left) and cell type (right) below. Cell type legend above Figure 5h. **j)** Barrett’s Esophagus tissue region E19_reg004 colored by expert annotated CN; color legend in colored boxes along x axis of Figure 5f. Extended view of black box region colored by CN (left) and cell type (right) below. Cell type legend above Figure 5h.

We applied this framework to a spatial proteomic dataset of Barrett’s Esophagus containing tissue samples from patients with heterogenous tissue morphology ranging from normal morphology to metaplasia, dysplasia, and tumor^49^ (**Fig. 5b**). This allows us to compare both disease stage (i.e., normal, metaplasia, dysplasia, or tumor) and individual (i.e., region) differences in MINGL assigned neighborhood probabilities. We also chose this dataset as each cell had been previously annotated with an assigned cellular neighborhood (CN) (**Fig. 5b**).

To understand changes in CN probabilities that are greatest with disease states, we calculated the delta between global and disease state specific MINGL probabilities. At a high level, this largely showed that there is a higher probability in the assignment of a cell to a specific CN in the same sample compared to when all cells (global) are used (**Supplemental Fig. 4a**). These decreases in the probability of CN assignment reflect additional heterogeneity in cell neighborhood composition when additional samples are in consideration. Specifically, comparing disease states we observed that the tumor disease state had the highest delta in probabilities (**Fig. 5c**). This change reflects that CN spatial compositions are most different in tumor state from the other conditions, even across compartmentalized CNs.

We also investigated these changes by evaluating the Shannon entropy of each cellular neighborhood within each state. While Shannon entropy doesn’t measure variance, it does measure the diversity and ratio of a multi-component system allowing us an orthogonal method to observe global changes in composition. We compared the Shannon entropy of the cell type composition per CN across disease states. We observed minor increases in the average diversity in diseased tissues and consistently large variance between different CNs across conditions (**Supplemental Fig. 4b**).

More specifically, the *Mature Intestinal and Immune* CN had the highest entropy across disease states (**Supplemental Fig. 4c**) and the largest change in assigned CN probability, particularly in the tumor state (**Fig. 5c**). This neighborhood is enriched with immune, stromal, and epithelial cell types in contrast to some of these other neighborhoods which contain largely one major cell type grouping (**Supplemental Fig. 4d**). Thus, this represents an important neighborhood at the interface of inflammation and invasion and large changes in composition in the tumor samples highlight this neighborhood as one to focus on.

Evaluating which cell types differ most in composition of the *Mature Intestinal and Immune* CN revealed that the largest differences came from changes in epithelial subtypes (**Supplemental Fig. 4e**). In tumor classified samples we see increases in cell type proportion of epithelial MUC1+ Ki67+ cells and decreases in epithelial CK7+ cells (**Fig. 5d**). This aligns with current understanding of a switch from a more metaplastic-like epithelial (CK7+) to a more aggressive (Ki67+) and stressed (MUC1+) epithelial cell found in dysplasia and cancer. Moreover, in the stroma there are increases in exhausted PD1+ CD4+ T cells and neutrophils and a loss of CD36+ endothelial and nerve cells, suggesting increased levels of inflammation (**Supplemental Fig. 4e, Fig. 5d**). The fact that other cell type proportions remain relatively constant and that this neighborhood proportion is similar across disease states (**Supplemental Fig. 4f**) suggests that these cell types could play important roles in the conversion of epithelial cells to a more aggressive state.

In contrast, plasma cells in the *Plasma Cell Enriched* neighborhood had some of the smallest changes in probabilities, indicating less heterogeneity in this CN (**Fig. 5c**). Even in the tumor disease state, which had the highest summed deltas across all CNs, the proportion of plasma cells in this neighborhood remained consistent (**Fig. 5e, Supplemental Fig. 4g**). The *Plasma Cell Enriched* CN is a cell neighborhood that arose upon metaplasia and continued through disease states and was found to be associated with better outcomes in disease progression^64^. It also resembles a similar *Plasma Cell Enriched* CN from the healthy human intestine which is also very compartmentalized and was shown to be enriched for specific cell-cell interactions to maintain a localized environment^11^. This indicates that structures containing dense plasma cells with other immune cells (e.g., CD4+ T cells, macrophages) are highly conserved and these cell type compositions are required for maintenance and do not vary much even with disease state.

We next used MINGL to define individual region and patient-specific differences in cell type composition of CNs (**Fig. 5f**). Region E11_reg006 from patient E11 has the largest summed delta probabilities across CNs, indicating that its spatial organization is the most different from global. Consistent with our prior findings at the disease state level, the *Mature Intestinal and Immune* CN had one of the greatest delta values and E11_reg006 is a tumor sample (**Fig. 5f**). Indeed, tumor samples often represent the greatest deltas from global probabilities and normal regions are the least deltas (**Fig. 5f**). Dysplasia and metaplasia regions span intermediate delta values. This suggests that as Barrett’s Esophagus progresses to tumor, conservation of compartmentalized cellular neighborhoods decreases.

To understand which patient has the most heterogeneity in their spatial organization, we summed the mean delta probabilities of each patient’s regions (**Supplemental Fig. 4h**). Patient E11 had the greatest summed change (**Supplemental Fig. 4h**), yet normalized by total number of regions, patient E19 had the greatest average change (**Supplemental Fig. 4i**). Interestingly, regions from the same patient, and sometimes also the same disease state, have diverse delta values (**Supplemental Fig. 4h**). This is notable because it highlights how particular spatial organizations may be heterogeneous across different tissues from the same patient and can point to particular patients that may have more divergent spatial biology than an average cohort.

Next, we sought to reveal specific cell types that are differentially enriched in the spatial organization of individual patients and specific regions. We focused our analysis on region E11_reg006, patient E11, and on the *APC Enriched Immune* CN for the high heterogeneity we had observed in this neighborhood and patient region (**Fig. 5f**). The *APC Enriched Immune* CN is globally characterized by high enrichment of professional antigen presentation cells (e.g., dendritic cells, M1 macrophages) and other immune and vasculature cell types (e.g., M2 macrophages, CD4+ and CD8+ T cells, lymphatic cells, endothelial cells) (**Supplemental Fig. 4d**).

Specifically for region E11_reg006’s *APC Enriched Immune* CN, the largest difference from global was a broad decrease in immune cells (e.g., CD8+ T cells, CD4+ Tregs, M2 macrophages, and CD4+ T cells) and the increase in epithelial pH2AX+ cells, squamous p63+ cells, and lymphatic cells (**Fig. 5h, Supplemental Fig. 4j**). There were still immune cells present in this neighborhood, but their numbers were reduced in E11_reg006 (**Fig. 5i**) relative to the global CN composition and to the same CN in another tumor region E19_reg004 (**Fig. 5j**). Importantly, despite these compositional shifts, M1 macrophages remained enriched in E11_reg006, with their log2 enrichment close to zero (**Fig. 5h**). This indicates that the defining anchor cells of the *APC Enriched Immune* CN are preserved, even as the broader immune context is altered.

Investigating the spatial organization of the M1 macrophage anchor cells further revealed heterogeneous patterns across tumor regions from different patients. In E11_reg006, M1 macrophage neighbors were observed to be epithelial pH2AX+ cells (**Fig. 5i**) whereas in E19_reg004 they were observed to be CD4+ T cells and CD8+ T cells (**Fig. 5j**). Thus, although the anchor cell identity of the *APC Enriched Immune* CN is maintained, differences in local associations between innate and adaptive immune arms exist in this CN across different patients.

Similarly, we observed changes in cell type proportion in the *APC Enriched Immune* CN for E11_reg006 compared with all E11’s samples (**Supplemental Fig. 4k**, **l**), demonstrating heterogeneity in spatial organization composition exists even within a single patient. These results indicate how M1 macrophages act like anchors of core functions of innate immunity to define the *APC Enriched Immune* CN but are less associated with other immune cells in patient E11 and to a greater extent in specific region E11_reg006 compared to global. This may indicate either an earlier or later state of the inflammation pathway. Finally, these results further highlight how MINGL can identify patient- and region-specific compositional changes in spatial architecture.

Together, these analyses demonstrate how MINGL can be used as a quantitative framework for measuring spatial heterogeneity in organizational structures. This can be done across any comparative group, such as patient, disease state, and tissue region. We used this to identify conserved and heterogeneous spatial architecture patterns across Barrett’s Esophagus progression and between patients and patient-specific regions. MINGL can thus identify patients with major differences in generally conserved tissue structures. This can lead to targeted analysis of most important organizational changes associated with a comparator like disease progression. Consequently, this is helpful to guide which organizational units and biological scales to focus on, regardless if they’re highly heterogeneous or uniform. These findings establish MINGL probability changes as a novel, versatile metric for defining spatial architecture heterogeneity.

### 2.5 MINGL Identifies Optimal Cluster Numbers for Biologically Relevant Cellular Neighborhoods

While our prior work with MINGL has focused on pre-existing cellular neighborhood (CN) labels, we also acknowledge that generating these CN labels is difficult, requiring guess-and-check workflows that can dramatically influence downstream organization associations. A major bottleneck and area of uncertainty in the spatial biology field is determining the optimal cluster number for defining CNs. This often results in the use of the default number of clusters by an unsupervised algorithm that may not be biologically most relevant. For those that explore cluster numbers further, selection is largely driven by human intuition, trial-and-error checking of multiple clusters through interacting with cluster compositions and alignment with prior literature. Due to the manual nature of these tasks, determination of the optimal cluster number for CNs from single-cell spatial data requires substantial time, expertise, and experience rather than quantitative measurements. Thus, there is a need for additional quantitative metrics to enable quick, unsupervised, and biologically relevant determination of the optimal cluster number for cellular neighborhoods of spatial-omics datasets.

Quantitative determination of the optimal cluster number for a given multidimensional dataset can be thought of as a balancing act of two competing forces: fit and confidence (**Fig. 6a**). On one hand, lower numbers of clusters will “separate” better than larger cluster numbers enhancing confidence in cluster identity (greater distance to neighboring cluster centroids), while on the other hand, higher numbers of clusters increase fit, as individual cells are more similar to their assigned clusters (shorter distance to centroid of assigned cluster). We hypothesized that we could leverage MINGL’s cluster assignment probabilities to quantify confidence and the per-neighborhood log-likelihood of feature similarity to the assigned cluster to characterize fit.

**Figure 6:**
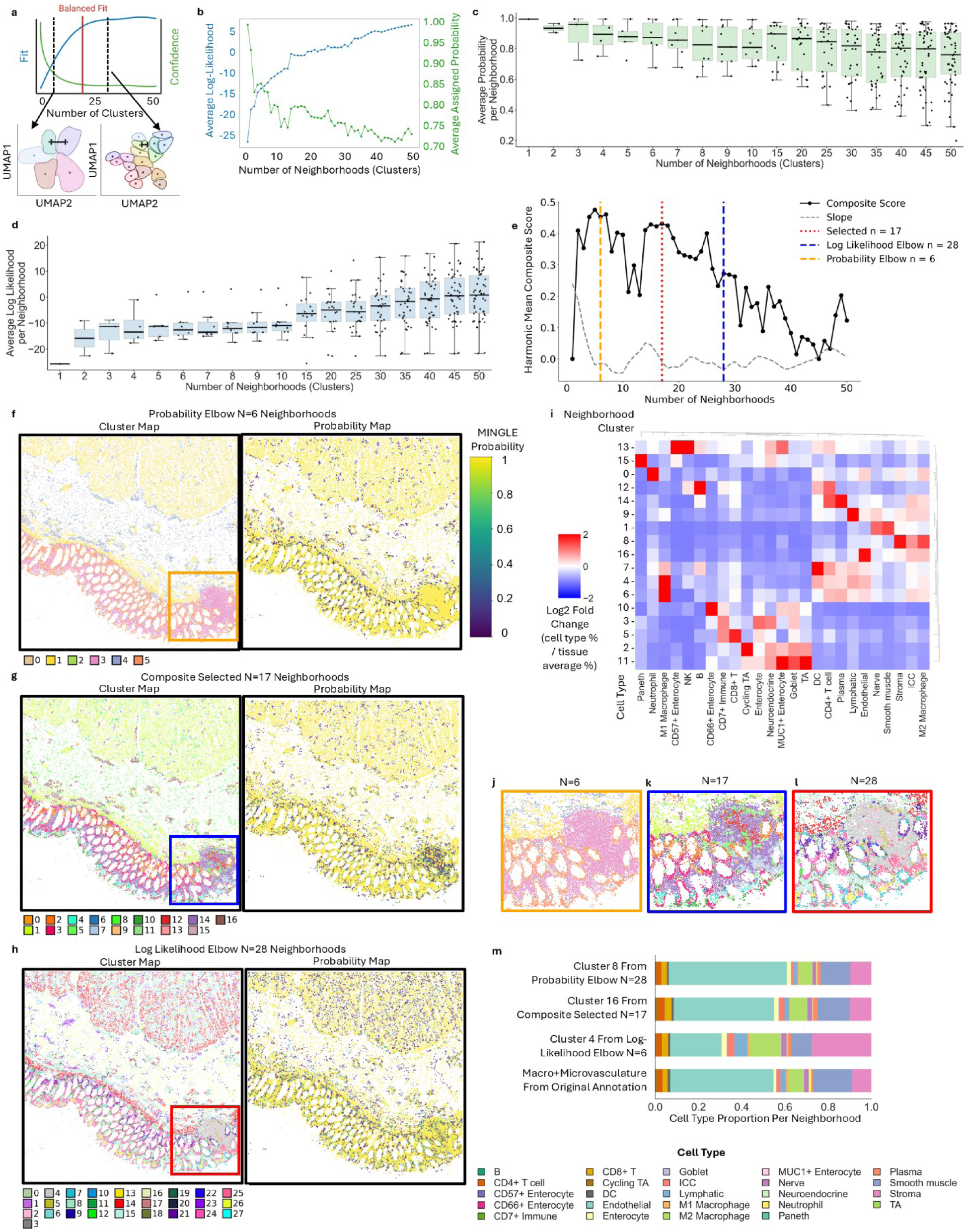
MINGL identifies cluster numbers for biologically relevant cellular neighborhoods. **a)** Tradeoffs between clustering fit and confidence as cluster number increases for number of cellular neighborhoods. **b)** Average log-likelihood and assigned cluster probability of all cells in the healthy intestine dataset from N=1 to 50 clusters for human intestine spatial dataset. **c)** Average probability per cluster distribution from N=1 to 50 clusters with box plots showing standard deviation and n=number of neighborhoods. **d)** Average log-likelihood per cluster distribution from N=1 to 50 clusters with box plots showing standard deviation and n=number of neighborhoods. **e)** Harmonic composite score (solid black) and composite score slope (dashed gray) from N=1 to 50 clusters. Calculated probability elbow (vertical dashed yellow), log-likelihood elbow (vertical dashed blue), and algorithm selection (vertical dashed red) cluster numbers are plotted. **f-h)** Representative healthy intestine region colored by **f)** probability elbow N=6 clusters, **g)** composite selected N=17 clusters, and **h)** log-likelihood elbow N=28 clusters (left) and assigned cluster probability (right). Color legend for clusters in Figure 6f-h. Legend for all probability plots in Figure 6f**. i)** Log2 fold change enrichment of cell types in each composite selected cluster (N=17). Black square region of an immune follicle from Figure 6f-h colored by **j)** probability elbow (N=6) clusters, **k)** composite selected (N=17) clusters, or **l)** log-likelihood elbow (N=28) clusters. **m)** Cell type proportions of vasculature clusters across different clustering numbers and from original expert annotations. Cell type color legend below Figure 6m.

We investigated MINGL assigned CN probability and log-likelihood values on the healthy human intestine dataset after K-means neighborhood clustering of single-cell neighbor windows^11^ using cluster numbers ranging from N=1 to 50. As expected, as the cluster number increased, the average log-likelihood across all cells in the dataset increased logarithmically and the average probability of a cell’s assigned cluster decreased in an exponential decay (**Fig. 6b**). Evaluating the variance in these distributions per-neighborhood, we observed that the variance of probability (**Fig. 6c**) and log-likelihood (**Fig. 6d**) also increased as cluster number increased. Applying the elbow point method^65^ for these metrics revealed dichotomous optimal cluster number suggestions of N=6 and N=28 CNs (**Fig. 6e**). Although both cluster numbers can be used to find interesting multicellular organizations, clustering at N=6 in this dataset lumps key distinct organizational structures of the intestine together (**Supplemental Figure 5a**), and N=28 produces good resolution on these structures but sometimes splits a CN that should only be one structure (**Supplemental Figure 5b**). Consequently, we hypothesized that these two metrics could offer biological bounds (two extremes of high confidence and low fit, or high fit and low confidence) and provide potential to create a composite metric that balances both for a more biologically relevant starting point for optimal CN number selection.

To create a quantitative composite metric from MINGL for determining CN numbers, we calculated a harmonic composite score combining log-likelihood and probability. We determined an optimal number of neighborhoods based on the composite score stability (i.e., slope of the composite score was near zero) within the cluster number range defined by the elbows of the probability and log-likelihood curves. The algorithm selected N=17 as the optimal cluster number for neighborhood clustering of the human intestine dataset (**Fig. 6e**). Our method improves upon classical approaches by capturing nuanced multi-neighborhood membership through probability-derived confidence and log-likelihood-derived fit, while also reducing guess-and-check workflows in neighborhood resolution selection, providing a biologically meaningful, quantitative starting point for selecting cluster numbers.

To understand the distribution of MINGL probabilities and biological relevance of the selected cluster number, we compared the localization and composition of the neighborhoods from each clustering resolution. Focusing in on just the mucosa region of the lowest clustering resolution (N=6) we see high MINGL probabilities, but the mucosa is largely only separated into epithelial and non-epithelial neighborhoods (**Fig. 6f**)—merging and missing key neighborhoods such as the *Plasma Cell Enriched* CN^11^. On the other hand, the composite selected resolution (N=17, **Fig. 6g**) and the log-likelihood elbow (N=28, **Fig. 6h**) resolve such structures in the mucosa yet have lower levels of CN probability assignment. However, clustering at N=28 resulted in two *Plasma Cell Enriched* CNs (15 and 18 of **Supplemental Fig. 5b**) that should be merged. In contrast, composite selected clustering at N=17 only has one *Plasma Cell Enriched* CN (14 of **Fig. 6i**).

Another example we can observe is the immune follicle, where at the lowest clustering resolution (N=6) the follicle structure is lumped together with the *Plasma Cell Enriched* CN (**Fig. 6j, Supplemental Fig. 5a**). On the other hand, at the composite selected (N=17) (**Fig. 6k**) and high clustering resolution (N=28) (**Fig. 6l**) we not only see separation from these other organizational structures, but we also observe separation of inner (i.e., N=17 #12—red, N=28 #2—light pink) and outer (i.e., N=17 #7—light blue, N=28 #4—light gray) follicle CNs. This corresponds to the lower assignment probabilities in the immune follicle as described earlier (**Fig. 1d**)—matching prior annotations reported^11^.

Finally, to investigate how neighborhood composition changes at different clustering resolutions, we compared the cell type proportions of endothelial enriched CNs at different cluster numbers (i.e., N=6, N=17, and N=28) to the combined *Microvasculature* and *Macrovasculature* CNs of the annotated dataset^11^. Cluster 4 from the lowest clustering resolution N=6 was the most different from the original annotations with a lower proportion of endothelial cells and a higher proportion of stroma and M2 macrophages (**Fig. 6m**). Cluster 16 from the composite selected resolution N=17 and Cluster 8 from the highest clustering resolution N=28 had similar cell type proportions to the original annotations and had a similarity score greater than 90% (**Supplemental Fig. 5d**). The results from these three vignettes suggest that at low clustering resolution, while assignment probability is high, meaningful spatial organization at the resolution of the data is lost. Conversely, the addition of more clusters can resolve all cellular neighborhoods but artificially splits several that should be clustered together based on cell type composition. It further emphasizes that MINGL’s quantitatively selected optimal cluster number has agreement with previously annotated CNs and can be used as a good starting point for neighborhood analysis.

Together, these results demonstrate that MINGL provides a quantitative framework for identifying cluster numbers for neighborhood analysis in spatial-omics that balances clustering fit and confidence while maintaining biological interpretability and relevance. By integrating log-likelihood and probability in a composite score that rewards stability, MINGL defines a biologically meaningful number in clustering resolution that provides a relevant starting point for spatial organization analysis. MINGL’s quantitatively selected optimal cluster number and edge conditions showed agreement with manual expert annotations and classical clustering performance indexes, further highlighting its utility for spatial-omics analyses while substantially reducing the time and expertise required for manual cluster number optimization.

## 3 Discussion

A central challenge in spatial biology is reconciling the inherently continuous nature of tissue organization with the discrete abstractions required for quantitative analysis. Although recent spatial-omics methods have enabled increasingly detailed identification of cellular neighborhoods and organizational structures, they largely treat organization as a set of internally homogeneous units with fixed boundaries^10^. This framing obscures biologically important features such as interfaces, gradients, and uncertainty—hallmarks of how tissues are assembled, maintained, and remodeled. Here, we present MINGL as a probabilistic framework that reframes tissue architecture as a landscape of graded organizational membership, enabling borders, transitions, and heterogeneity to be quantified across hierarchical biological scales.

A defining contribution of MINGL is the explicit identification and characterization of border cells—cells that reside at the intersection of multiple organizational units. Unlike existing approaches that treat such cells as noise or force them into a single assignment, MINGL leverages probabilistic membership to reveal borders as conserved and biologically meaningful regions of cellular mixing. Across both diseased (melanoma) and healthy (human intestine) tissues, border cells were consistently enriched at known anatomical and functional interfaces and exhibited distinct cell-type enrichments not apparent from neighborhood composition alone.

In particular, we observed enrichment of stromal, dendritic, and macrophage populations at the tumor border, while in the healthy intestine, dendritic cells and macrophages were similarly enriched at the mucosa–muscularis mucosa interface. These results suggest conserved roles for innate immune cells in regulating epithelial–stromal transitions^66,67^ and should be studied further as they could be therapeutically targeted for processes such as tumor invasion. These results support the view that borders are not merely geometric boundaries, but active regulatory zones enriched for specific cell types that may coordinate signaling, communication, and structural integrity between adjacent organizational units. By making borders quantitatively accessible, MINGL extends existing neighborhood-based approaches toward a more complete description of tissue organization.

Further, MINGL uses identification of these borders to construct interaction networks of higher-order tissue organization. This contrasts with most spatial analyses that solely focus on identification of cellular neighborhoods or a single scale, not how they are also spatially organized and relationships across scales. Understanding how higher-order organizational structures interact is essential for explaining how local cellular coordination scales to tissue-level function.

At tissue unit, community, and cellular neighborhood scales, the MINGL interaction networks recapitulated known spatial hierarchies—such as the layered organization of the intestine—while simultaneously exposing hubs of interaction that span epithelial, stromal, and immune compartments. Notably, plasma cell–enriched communities and neighborhoods emerged as highly connected nodes across multiple scales. Although plasma cells are not traditionally viewed as structural organizers, their conserved localization to specific interfaces suggests a potential architectural role, for example by balancing long-term immune residence with access to epithelial transport pathways^59,68^. These observations illustrate how MINGL-derived interaction graphs can reveal candidate organizing units and cross-compartment coordinators.

A second conceptual advance of MINGL is the ability to quantify not only where organizational units meet, but how they transition. MINGL complements current expression-based gradient methods^24,51,69,70^ by revealing cell types and organizational changes associated with molecular gradients at borders. By using neighborhood assignment probabilities, MINGL can quantitatively capture transitions spanning sharp borders to gradual gradients across biological scales to highlight critical cellular interactions.

Applied to immune follicles, this approach recapitulated known center–periphery organization while revealing continuous redistributions of cell types within neighborhoods previously treated as uniform. Extending this analysis to another epithelial and immune cellular neighborhood interface in the intestine had a more gradual transition, demonstrating that different organizational interfaces are governed by distinct transition dynamics. This heterogeneity suggests that gradient steepness itself may be a regulated feature of tissue architecture and another dimension on which to evaluate organization.

With MINGL’s multimember neighborhood assignments, we also created a general framework for comparing organizational structures across conditions. Unlike differential abundance analyses, which collapse subtle compositional changes, this approach captures condition-associated reorganization driven by shifts in a limited number of cell types that are obscured by discrete clustering. These probability-based metrics introduce an orthogonal axis of comparison that directly reflects architectural variation rather than changes in cell neighborhood prevalence alone.

Applied to Barrett’s Esophagus progression, this framework identified neighborhoods that were selectively reorganized with disease, alongside others—such as the plasma cell niche—that remained remarkably stable. Consistent with our findings in the intestine, this underscores the strong compartmentalization of plasma cell niches across organs and disease states. More broadly, MINGL reveals heterogeneity in which cellular neighborhoods are altered across samples, enabling identification of disease-relevant organizational changes, stratification of patients, and detection of outliers. Together, these results suggest that architectural heterogeneity itself constitutes a biologically informative phenotype.

MINGL also addresses a persistent challenge in spatial analysis: selecting an appropriate number of neighborhoods. Rather than relying on manual curation or single-metric optimization, MINGL integrates assignment confidence and clustering fit into a composite score that identifies biologically meaningful bounds and a stable resolution range. This approach does not prescribe a single “correct” number of neighborhoods but provides an unbiased, biology-guided range that balances over-merging and over-splitting of organizational units. The resulting neighborhood structures show strong agreement with expert annotations while reducing user-dependent bias and analysis time, an advantage that will become increasingly important as spatial datasets grow in size and complexity.

We designed MINGL to be practically usable and broadly applicable. It is implemented as a scverse python package and has options for both CPU and GPU accelerated analysis enabling analysis with low compute across large single-cell spatial datasets. Moreover, because MINGL takes as input cell type labels, this enables it to be modality agnostic and be applicable to all spatial-omics datasets. Finally, while we demonstrated MINGL in the use case of single-cell spatial-omics, the underlying framework of MINGL can be applied to any spatial coordinate data with categorical values, which could enable its application to datasets in the fields of ecology, meteorology, and public health.

Taken together, MINGL advances spatial analysis by unifying discrete and continuous representations of tissue organization within a single probabilistic framework. In contrast to methods that focus solely on defining neighborhoods, spatial correlations, or interaction frequencies, MINGL treats uncertainty, mixing, and hierarchy as intrinsic properties of tissue architecture. This perspective enables borders, gradients, and heterogeneity to be analyzed using shared mathematical principles across biological scales and experimental contexts, extending beyond the capabilities of hard-clustering paradigms.

Looking forward, the MINGL framework opens several avenues for future investigation. MINGL is generally applicable across spatial-omics modalities, as it operates on cell-type-level annotations and can be applied to user-defined cellular neighborhood annotations generated by any upstream algorithm. Applied to longitudinal, developmental, or perturbation datasets, MINGL could be used to track how borders and gradients emerge, shift, or dissolve over time, providing insight into the dynamics of tissue assembly and remodeling. Integration with spatial transcriptomic, epigenomic, or imaging-based measurements could link architectural transitions to underlying molecular programs. In disease contexts, quantitative metrics of border composition, gradient steepness, or architectural heterogeneity may serve as biomarkers of progression or therapeutic response. Together, MINGL positions tissue architecture and spatial organization as a measurable, comparable, and actionable dimension of tissue biology.

## 4 Methods

### 4.1 Dataset Collection and Requirements

MINGL is implemented as a Python package for defining continuous spatial organization membership probabilities from discrete hierarchical annotations. The framework is compatible with single-cell spatial-omics datasets annotated at minimum with i) spatial coordinates, ii) cell-type labels, and iii) discrete hierarchical spatial organization labels (e.g., cellular neighborhoods, communities, or tissue units).

We applied MINGL to three previously published and peer-reviewed CODEX spatial proteomics datasets spanning healthy human intestine^11^, human melanoma^12^, and Barrett’s Esophagus disease progression^64^. All datasets were analyzed independently. Spatial coordinates and cell annotations were used as provided in the original publications without modification.

All analyses were implemented using the AnnData data structure (scverse ecosystem), enabling unified storage of per-cell metadata, neighborhood features, centroid parameters, and probability matrices.

### 4.2 Nearest Neighbor Feature Construction

For each cell, we constructed a vector of neighbor cell type composition using a k-nearest neighbors (kNN) approach^11,34,49^. Windows were computed independently within each tissue region to prevent cross-region mixing.

For every index cell, the *k* nearest neighboring cells were identified using Euclidean distance in two-dimensional spatial coordinate space. The index cell itself was excluded. Within each window, we counted the number of each lower-level hierarchical label (e.g., cell types, cellular neighborhoods). These counts form a feature vector describing the local microenvironment of that cell.

At the cellular neighborhood (CN) level, we used *k* = 10 nearest neighbors with cell type label features. At the community level, *k* = 100 with CN label features. At the tissue unit level, *k* = 300 with community label features. These values were selected based on prior hierarchical spatial organization analyses^11^ and reflect increasing spatial scales of organization.

To reduce boundary artifacts, neighboring cells located beyond a specified maximum spatial radius were excluded from windows.

### 4.3 Defining Centroids of Spatial Organization Units

MINGL models each annotated spatial organization unit as a centroid defined by the mean and spread of adjacent lower-level compositional features.

Cellular Neighborhood Centroids

For each expert-annotated cellular neighborhood 𝑛, we collected all cells assigned to that neighborhood. For each feature 𝑡 (e.g., a cell-type count within the kNN window), we computed the sample mean and sample standard deviation across cells in neighborhood 𝑛. These statistics define a univariate Gaussian distribution for each feature within each neighborhood. The full set of per-feature means and standard deviations constitutes the centroid representation of neighborhood 𝑛.

Higher Hierarchical Level Centroids

Centroids at higher hierarchical levels were constructed analogously. Community-level centroids were defined using nearest-neighbor cellular neighborhood composition features, and tissue unit centroids were defined using nearest-neighbor community composition features. Each hierarchical level was computed independently.

### 4.4 Gaussian Mixture Model Probability Assignment

To quantify graded membership of each cell to all centroids, MINGL uses a Gaussian mixture model framework with diagonal covariance structure. Each compositional feature is modeled as an independent univariate Gaussian, and features are assumed conditionally independent given the centroid.

For each cell, the likelihood of its observed feature vector under each centroid is computed as the product of per-feature Gaussian densities. Likelihoods are evaluated in log space for numerical stability and converted to normalized probabilities using a softmax transformation.

To prevent instability due to extremely small or zero variances, each standard deviation parameter is regularized using a minimum positive floor (ε = 10⁻⁶). The resulting per-cell probability vector defines a continuous representation of hierarchical spatial membership.

### 4.5 Identification of Border Cells

A probability threshold of ≥ 0.25 was used to define positive membership in a centroid. Cells exceeding this threshold in more than one centroid were classified as border cells.

For each cell 𝑖, we define the number of positive memberships as:

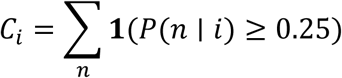

where 𝟏(⋅)denotes the indicator function. Cells with 𝐶_𝑖_ > 1 were considered border cells. The identities of co-positive centroids were recorded for downstream interaction analysis.

To assess robustness of border definitions, we repeated border identification across thresholds ranging from 0 to 0.25. At each threshold, border cell compositions and their spatial locations were recomputed and evaluated.

#### Border-Specific Cell-Type Enrichment

For a selected pair of centroids 𝐶𝑁1 and 𝐶𝑁2, we compared the cellular composition of cells positive for both 𝐶𝑁1 and 𝐶𝑁2 (border group) and cells positive for only one centroid (single groups).

For each cell type 𝑡, enrichment was defined as:

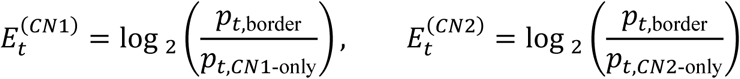

A cell type was considered enriched at the interface if enrichment was positive relative to both single-neighborhood groups.

### 4.6 Network Graph Construction

To characterize higher-order spatial interactions, we constructed weighted graphs of centroid co-membership. For cells characterized as a border cell, pairwise combinations of positive centroid identities are recorded. Edge weights between centroids correspond to the number of cells exhibiting co-membership.

#### Geometric Mean Network Graph Edge Weights

A complementary approach for defining neighborhood associations was quantified using identified border cells with at least two neighborhood probabilities exceeding 0.25. For each pair of neighborhoods 𝐶𝑁1 and 𝐶𝑁2, directional co-occurrence probabilities were computed as 𝑃(𝐶𝑁2 ∣ 𝐶𝑁1)and 𝑃(𝐶𝑁1 ∣ 𝐶𝑁2). Edge weights were defined as the geometric mean of these bidirectional probabilities, 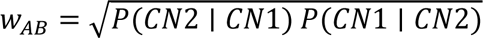, providing a symmetric measure of reciprocal neighborhood association. Edge thickness in network visualizations was scaled linearly to 𝑤_𝐴𝐵_.

### 4.7 Gradient-Based Quantification of Spatial Transitions

#### Probability-Weighted Log-Ratio Score

To quantify continuous transitions between two spatial organization units (e.g*., Inner Follicle* and *Outer Follicle* CNs), we defined a probability-weighted log-ratio score for each cell.

To quantify continuous transitions between two spatial organization units, we defined a probability-weighted log-ratio score. For neighborhoods 𝐶𝑁1and 𝐶𝑁2:

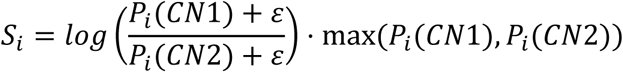

where 𝜀 = 10^−9^. This formulation captures both directional bias and assignment confidence.

#### Equal-Width Stratification of Probability Ratio Scores

Continuous scores were partitioned into ordered bins using equal-width discretization. For 𝐾bins:

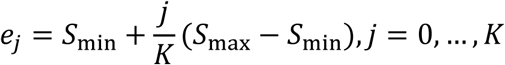

Cells were assigned to bins based on interval membership. Bins were labeled in ascending order (for example, 20th percentile through 100th percentile when 𝐾 = 5). This deterministic discretization preserves geometric spacing of the score distribution.

#### Deterministic Ordering of Transition Clusters

Clusters were ranked using a size-normalized weighted percentile score:

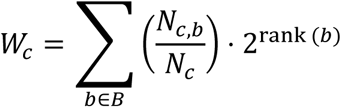

where 𝑁_𝑐,𝑏_is the number of cells in cluster 𝑐assigned to bin 𝑏, and 𝑁_𝑐_is the cluster size. Clusters were ordered by decreasing 𝑊_𝑐_.

#### Discrete Transition Derivatives and Steepness Score

To quantify the sharpness of transitions between ordered clusters, we computed first and second discrete derivatives of mean cluster scores.

Let 𝑆_𝑘_ denote the mean score of ordered cluster 𝑘.

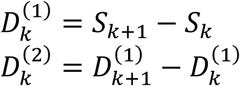

Steepness was defined as:

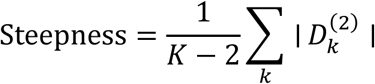

The steepness score metric quantifies curvature in the ordered transition sequence and provides a global measure of abrupt versus gradual structural change

Maximum curvature was also computed:

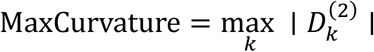

#### Local Spatial Gradient Estimation

To measure spatial transition dynamics directly within tissue coordinates, we estimated local score gradients using spatially restricted linear regression.

For each cell 𝑖, we identified its 𝑘 = 20 nearest spatial neighbors within the same region. We then fit a local linear regression:

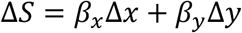

Gradient magnitude was computed as:

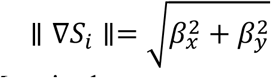

#### Spatial Normalization of Gradient Magnitude

Because gradient magnitude depends on the spatial scale of sampling, we normalized gradients by the characteristic neighbor spacing of each region.

Let 𝑑_NN_denote the mean nearest-neighbor distance within a region. Spatially normalized gradient magnitude was defined as:

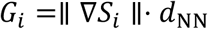

This normalization yields the expected change in score across a typical intercellular distance, enabling comparison of transition steepness across regions with differing cell densities.

Normalized gradients were visualized spatially and summarized using median and mean values within each region.

### 4.8 Defining Optimal Number of Neighborhoods

To identify the optimal number of unsupervised neighborhood clusters, we performed k-means clustering across a range of cluster numbers. For each 𝑛, we computed the average log-likelihood and average assigned probability under the Gaussian model.

Both metrics were normalized to [0,1]. We then defined a composite stability score using the harmonic mean:

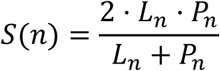

where 𝐿_𝑛_ is normalized average log-likelihood and 𝑃_𝑛_ is normalized average assigned probability for cluster number 𝑛.

Plateaus in 𝑆(𝑛) characterized by high score and low slope were identified as stable model regimes. The selected cluster number corresponded to the earliest stable plateau between elbow points.

### 4.9 Quantifying Context-Dependent Heterogeneity in Spatial Architecture

To quantify heterogeneity in multicellular organization across conditions (e.g., disease state, patient, anatomical region) we defined a metric for quantification of organizational tissue heterogeneity. This framework enables detection of spatial reorganization even when defining anchor cells remain consistent.

#### Context-Specific Spatial Heterogeneity (Δ-probability)

To measure how spatial organization changes across contexts, we first computed global MINGL probabilities using all cells in the dataset.

Next, the dataset was partitioned into context-specific subsets (e.g., disease state, unique tissue region). For each subset, centroids were recomputed independently, and new context-specific probabilities were calculated.

For each cell 𝑖 and centroid 𝑛, we defined the delta probability:

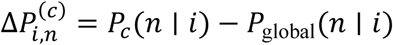

where: 𝑃_𝑐_(𝑛 ∣ 𝑖) is the probability computed using centroids derived only from context 𝑐, and 𝑃_global_(𝑛 ∣ 𝑖) is the probability computed using centroids derived from the full dataset.

Positive values indicate that, within context 𝑐, the cell is more strongly compatible with centroid 𝑛 than under the global model. Negative values indicate increased similarity to other centroids relative to global.

For each centroid and context, we computed the mean delta probability:

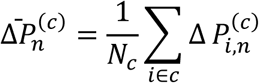

where 𝑁_𝑐_is the number of cells in context 𝑐. Total Heterogeneity Score for each Context
To identify which contexts (e.g., disease state, individual patient, unique tissue region) diverge most strongly from the global tissue architecture, we defined a global heterogeneity index for each context as the sum of absolute mean delta values across all centroids:

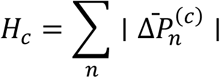

This metric quantifies total structural deviation from the global spatial model. Contexts with larger 𝐻_𝑐_values exhibit greater reorganization of multicellular architecture.

This index was used to identify disease states, patients, or regions that are most structurally distinct from the cohort average.

#### Patient-Level Aggregation of Heterogeneity

To quantify heterogeneity at the patient level, regional heterogeneity indices were aggregated per patient:

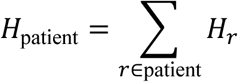

where 𝐻_𝑟_is the heterogeneity score of region 𝑟.

Patient-level heterogeneity was evaluated both as total summed deviation and normalized by the number of regions per patient to distinguish widespread versus localized divergence.

#### Context-Specific Compositional Enrichment or Depletion

Structural heterogeneity does not necessarily imply changes in the defining anchor cell type of a neighborhood. To assess compositional shifts within neighborhoods, we computed cell-type enrichment relative to the global reference.

For a given cell type 𝑡, neighborhood 𝑛, and context 𝑐, we defined enrichment as:

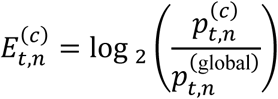

where 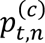 is the proportion of cell type 𝑡 within neighborhood 𝑛 in context 𝑐, and 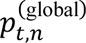 is the global proportion of cell type 𝑡 within neighborhood 𝑛.

Positive values indicate overrepresentation relative to global composition; negative values indicate depletion.

Only cell types exceeding 1% abundance in the context and exhibiting at least two-fold change (∣ 𝐸 ∣≥ 1) were annotated in enrichment plots.

Context-specific abundance was visualized using size-scaled dot plots. Dot size was proportional to the percentage abundance of a cell type within a context. Larger dots with larger fold change magnitude are most different from global composition.

#### Shannon Entropy of Neighborhood Composition

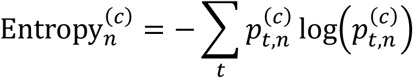

Entropy was used as an orthogonal measure of compositional diversity within neighborhoods.

## 5 Data Availability

The CODEX healthy human intestine dataset used in this study can be found in the online repository at Dryad https://doi.org/10.5061/dryad.pk0p2ngrf

The CODEX human melanoma dataset used in this study can be found in the online repository at Dryad https://doi.org/10.5061/dryad.k0p2ngfcc

The CODEX Barrett’s Esophagus disease progression dataset can be found in the online repository at Zenodo https://doi.org/10.5281/zenodo.14976466

## 6 Code Availability

MINGL was written in Python as a PIP installable package. Source code, tutorial notebooks, and documentation are available on Github at https://github.com/HickeyLab/Mingl.

## 7 Acknowledgements

This work was supported by the US National Institutes of Health (grant nos. 3U54AG07593, 3OT2OD033759-01S4, 3OT2OD033759-01S5, 1U01-AI186999-01, 1U54-AI191253-01); the National Institute of General Medical Sciences NIH National Service Research Awards Program T32 Training Grant (1T32GM144291); the National Science Foundation CAREER Award (grant no. 2440733); the V Foundation (V2025-019); the Human Frontier Science Program Early Career Grant (RGEC26/2025-); and start-up support from Duke University Department of Biomedical Engineering for JWH.

