## Supplemental Information for "MINGL Quantifies Borders, Gradients, and Heterogeneity in Multicellular Tissue Organization"

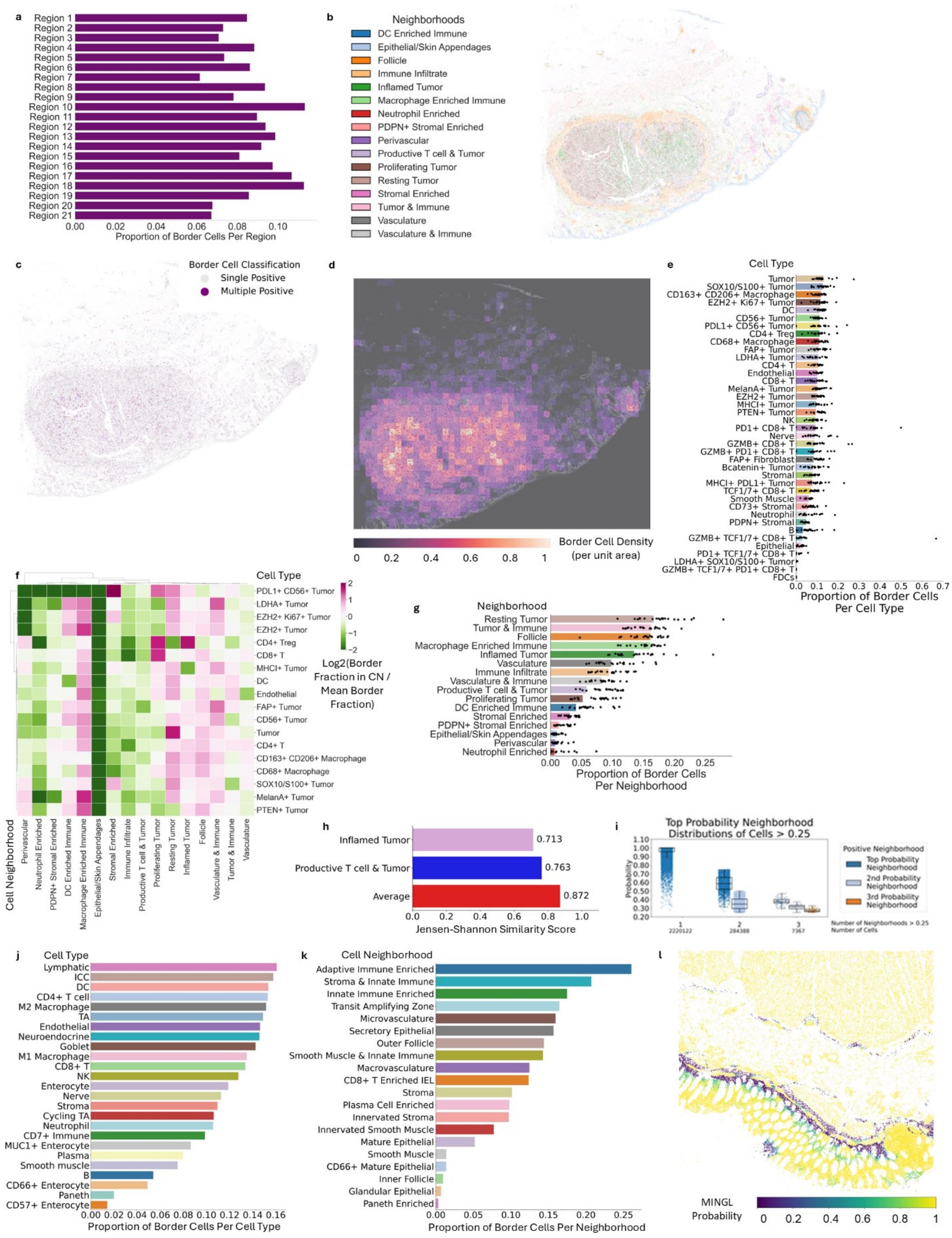

**Supplemental Figure 1: Border Cells in Melanoma and Healthy Intestine a)** Proportion of border cells per unique region in human melanoma. **b)** Spatial location of cellular neighborhoods

(CN) in a representative melanoma tissue region. Color corresponds to CN in Neighborhood legend. **c)** Spatial location of multiply positive border cells (purple) and singularly positive non-border cells (gray) in the representative melanoma region from **b. d)** Border cell density per unit area in representative melanoma region from **b** and **c. e)** Proportion of each cell type classified as a border cell in melanoma. Replicates per unique tissue region are shown in black dots. **f)** Log2 fold enrichment of border cells in each CN. Enrichment calculated as the fraction of border cells among all cells of a given cell type assigned to that CN, normalized to the mean border-cell fraction for that cell type across all CNs. **g)** Proportion of each CN classified as a border cell in melanoma. Replicates per unique region are shown in black dots. **h)** Jensen-Shannon similarity score of border cell type composition with *Inflamed Tumor*, *Productive T Cell & Tumor*, or average of *Inflamed Tumor* and *Productive T Cell & Tumor* CNs. **i)** MINGL defined CN membership probability distributions of healthy human intestine cells. Threshold for positive assignment is 0.25. **j)** Proportion of each cell type in healthy human intestine dataset classified as a CN border cell. **k)** Proportion of each CN in healthy human intestine dataset classified as a CN border cell. **l)** MINGL probability of expert annotated tissue unit label in a representative healthy intestine region.

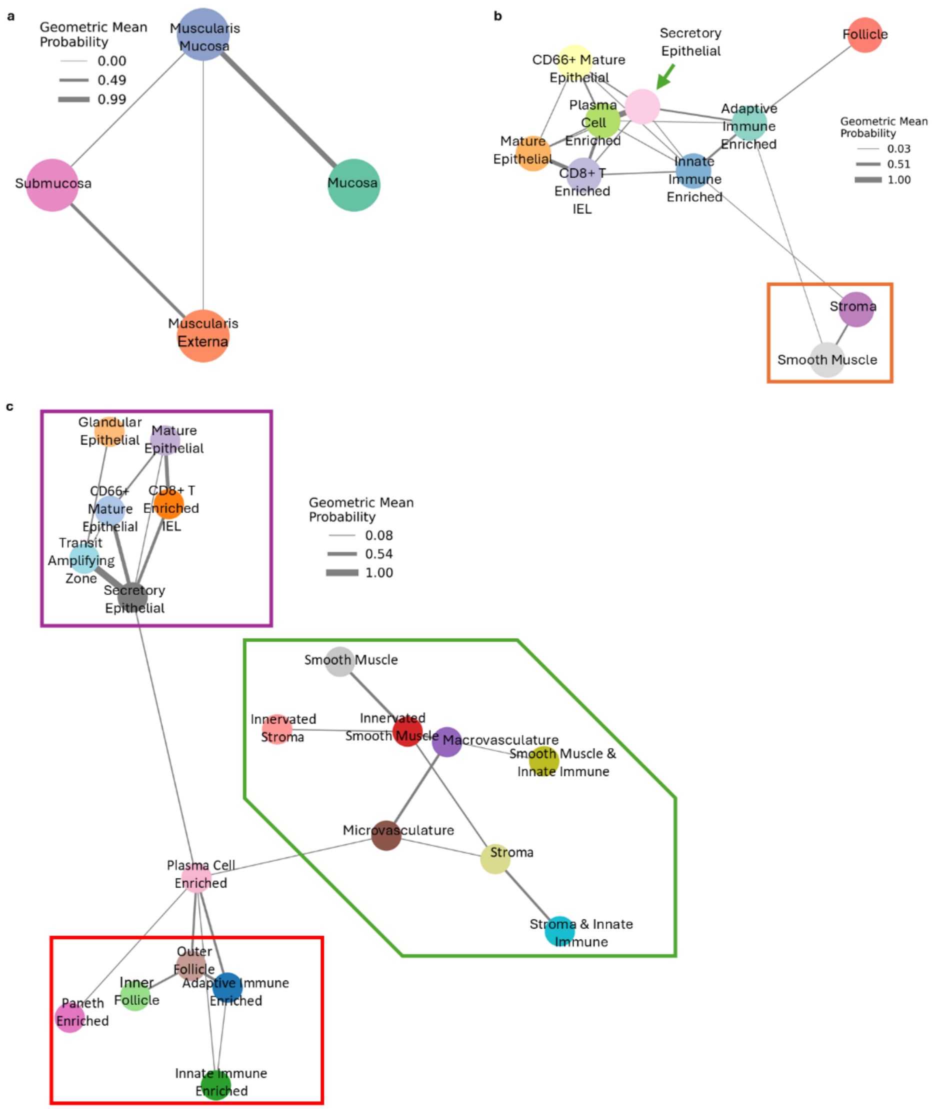


**Supplemental Figure 2: Geometric Mean Probability Hierarchical Spatial Organization Network Graphs a)** Top 4 tissue unit interactions network graph. Edge thickness represents the geometric mean of the bidirectional co-occurrence probabilities between each pair of neighborhoods, calculated from the proportion of cells in one cell neighborhood (CN) also positive for another CN. **b)** Top 20 community interactions network graph. Edge thickness between nodes is geometric mean described in **a**. **c)** Top 25 CN interactions network graph. Edge thickness between nodes is geometric mean described in **a**.

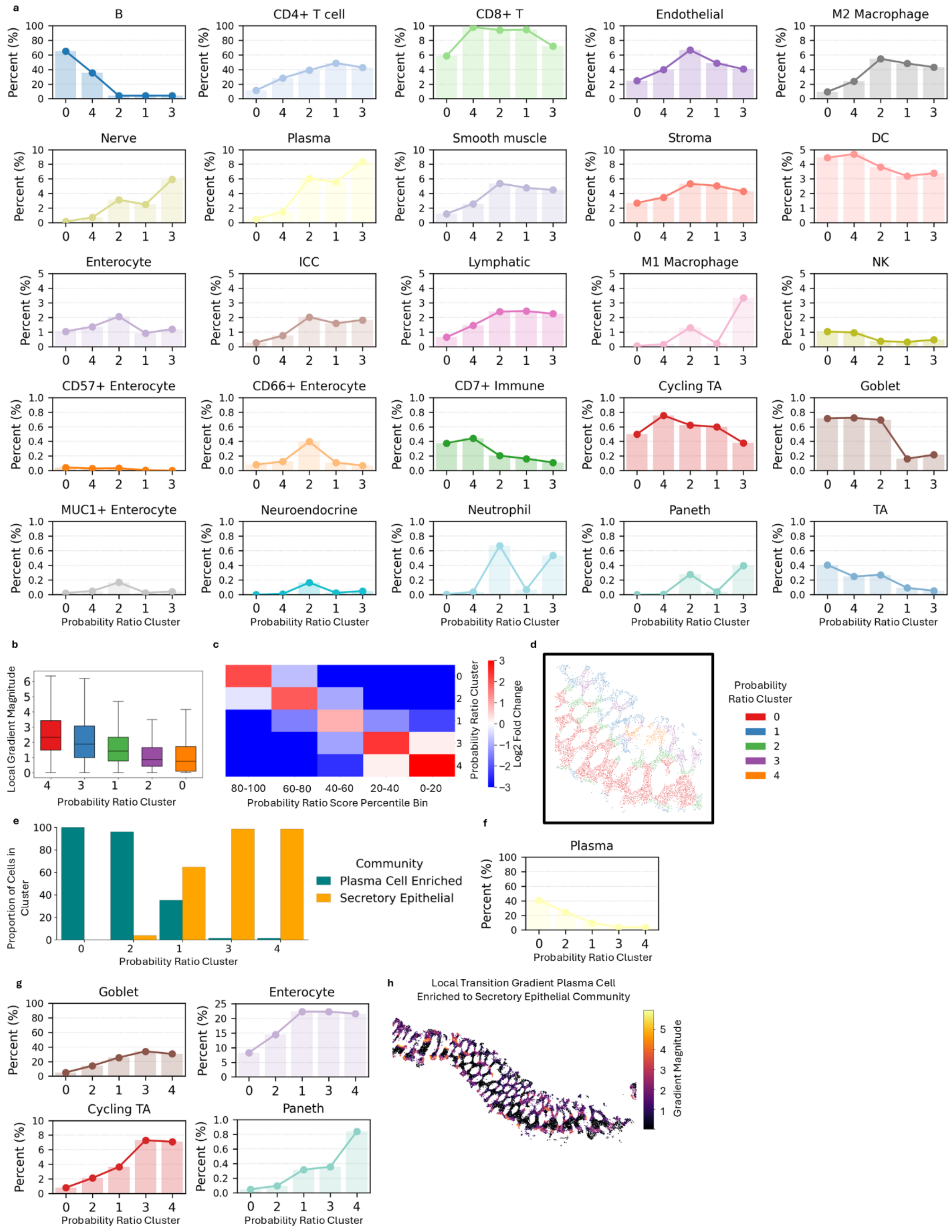


**Supplemental Figure 3: Spatial Gradients in Cellular Organization and Cell Type Composition a)** Expert annotated cell type proportion changes across ordered clusters between

*Inner Follicle* and *Outer Follicle* cell neighborhoods (CNs). **b)** Average local gradient magnitude of individual cells in each ordered cluster. Mean and standard deviation are shown. **c)** Log2 fold enrichment of probability ratio percentile across clusters for *Plasma Cell Enriched* and *Secretory Epithelial* communities in healthy human intestine. **d)** Expanded view of healthy intestine crypts from **Figure 5j.** Color corresponds to cluster from **c. e)** Proportion of *Plasma Cell Enriched* and *Secretory Epithelial* communities in each cluster from **c.** Clusters are ordered in descending order of highest enrichment of higher probability ratio percentiles. **f)** Expert annotated cell type proportion changes across ordered clusters from **c. h)** Local gradient magnitudes between the *Plasma Cell Enriched* and *Secretory Epithelial* communities of a healthy intestine.

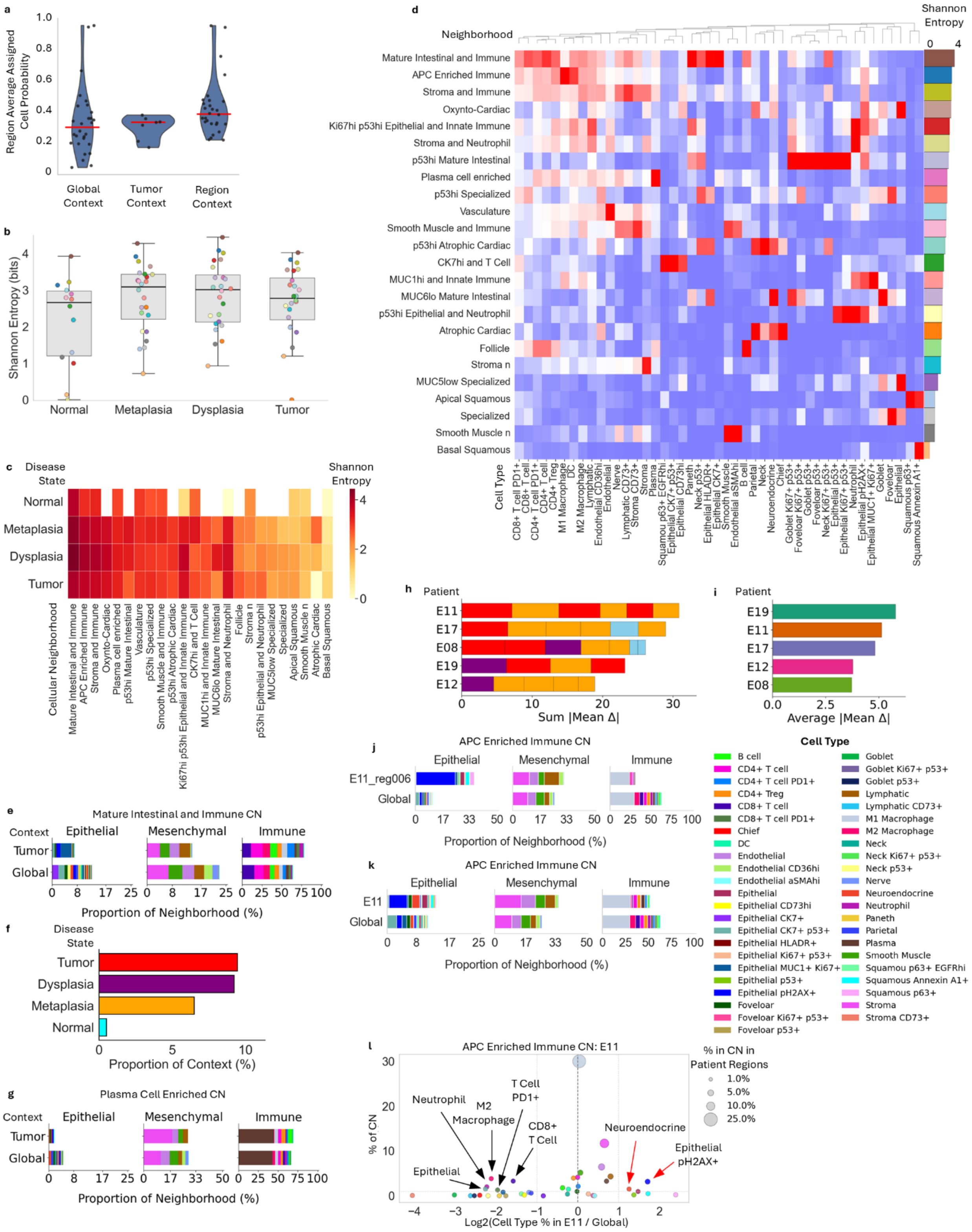


**Supplemental Figure 4: Spatial Heterogeneity and Diversity at Hierarchical Scales** All data from this figure is from the Barrett's Esophagus progression dataset. **a)** Average expert assigned cell neighborhood (CN) MINGL probability when MINGL is computed across groups (e.g. global or all regions, individual disease state, individual regions). Each dot represents the mean assigned probability of an individual region with the red bar as the mean across all regions. **b)** Shannon entropy of each disease state calculated as a measure of diversity of cell types within each CN. Dot color corresponds to CN in **d.** Shannon entropy calculated for each CN in each disease state. Mean and standard deviation are shown. **c)** Shannon entropy of each CN in each disease state. Shannon entropy calculated as a measure of cell type diversity in each CN. White boxes have no cells annotated as that CN in that disease state. **d)** Log2 fold enrichment of cell types in each CN from expert annotations. Shannon entropy is shown for cell type diversity in each CN. CNs are arranged in descending order of Shannon entropy. **e)** Cell type percentages in the *Mature Intestinal and Immune* CN for the tumor disease state and global (all disease states). **f) g)** Cell type percentages in the *Plasma Cell Enriched* CN for the tumor disease state and global (all disease states). **h)** Sum of the absolute value of the mean delta scores for each patient. Bars are split into individual region contributions to the total sum. Bar color corresponds to disease state as colored in **f. i)** Average of the absolute value mean delta scores for each patient normalized per unique region. **j)** Cell type proportions of the *APC Enriched Immune* CN in unique region E11_reg006. **k)** Cell type proportions of the *APC Enriched Immune* CN across all unique regions of patient E11. Color in **e, g, j, k** corresponds to cell type in cell type legend to the right of **j** and **k.** Cell types are displayed split into epithelial, mesenchymal, and immune cell types. **l)** Log2 fold change in *APC Enriched Immune* CN cell type proportions in all tissue regions of patient E11 compared to global. Dot color corresponds to cell type and dot size corresponds to the proportion of each cell type in the specific tissue region.

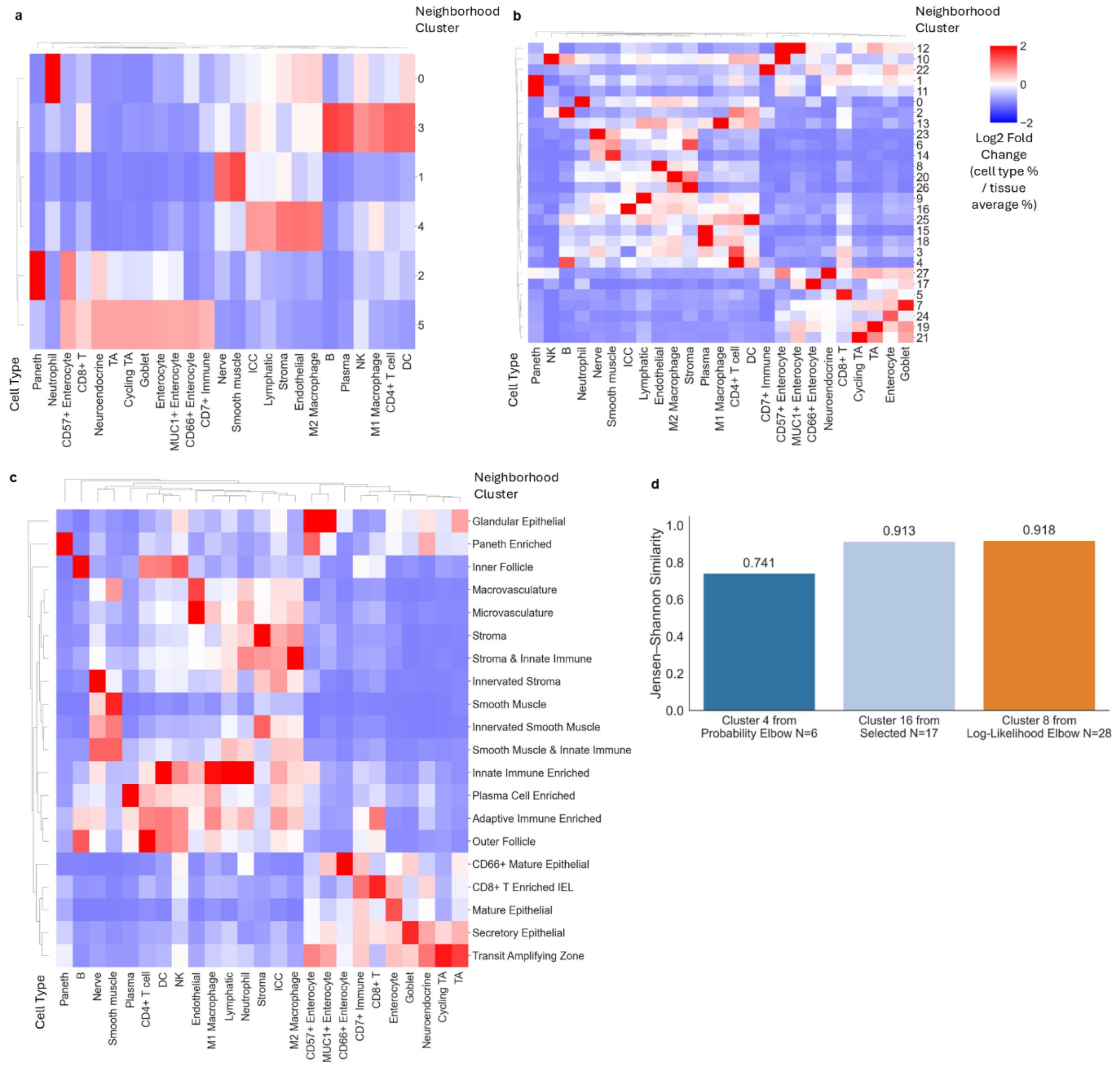


**Supplemental Figure 5: Cluster Cell Type Enrichment and Similarity a)** Log2 fold cell type enrichment of log-likelihood elbow N=6 clusters. Legend in **b. b)** Log2 fold cell type enrichment of probability elbow N=28 clusters. **c)** Log2 fold cell type enrichment of expert annotated N=20 clusters with expert annotated cluster labels. Legend in **b. d)** Jensen-Shannon similarity score of cell type proportions in elbows (N=6 and 28) and composite selected (N=17) cluster number compared to expert annotated vasculature clusters.
